# ATG8 delipidation is not universally critical for autophagy in plants

**DOI:** 10.1101/2023.08.23.554513

**Authors:** Yong Zou, Jonas A Ohlsson, Sanjana Holla, Igor Sabljić, Jia Xuan Leong, Florentine Ballhaus, Melanie Krebs, Karin Schumacher, Panagiotis N Moschou, Simon Stael, Suayb Üstün, Yasin Dagdas, Peter V Bozhkov, Elena A Minina

## Abstract

Intracellular recycling via autophagy is governed by post-translational modifications of the autophagy-related (ATG) proteins. One notable example is ATG4-dependent delipidation of ATG8, a process that plays critical but distinct roles in autophagosome formation in yeast and mammals. Here, we aimed to elucidate the specific contribution of this process to autophagosome formation in species representative of evolutionary distant green plant lineages: unicellular green alga *Chlamydomonas reinhardtii*, with a relatively simple set of *ATG* genes, and a vascular plant *Arabidopsis thaliana*, harboring expanded *ATG* gene families.

Remarkably, the more complex autophagy machinery of *Arabidopsis* rendered ATG8 delipidation entirely dispensable for the maturation of autophagosomes, autophagic flux and related stress tolerance; whereas autophagy in *Chlamydomonas* strictly depended on the ATG4-mediated delipidation of ATG8. Importantly, we uncovered the distinct impact of different Arabidopsis ATG8 orthologs on autophagosome formation, especially prevalent under nitrogen depletion, providing a new insight into potential drivers behind the expansion of the ATG8 family in higher plants.

Our findings underscore the evolutionary diversification of the molecular mechanism governing the maturation of autophagosomes in eukaryotic lineages and highlight how this conserved pathway is tailored to diverse organisms.

## Introduction

Autophagy is a catabolic pathway which plays a vital role in sustaining functionality of cellular constituents and in helping organisms overcome nutrient scarcity. Autophagy balances out the biosynthetic activity of cells and salvages nutrients by sequestering expendable or dysfunctional cytoplasmic content into specialized double membrane-vesicles, autophagosomes, and delivering them to the lytic compartment for degradation and recycling^1^. Effective cargo sequestration relies on continuous expansion of the autophagosomal membrane to create adequately sized vesicles. Subsequently, these autophagosomes must be sealed before fusing with the lytic compartment to avoid the leakage of hydrolases into the cytoplasm^2^.

The process of autophagosome formation and maturation is orchestrated by the autophagy-related proteins (ATGs) conserved among eukaryotes^1,3^. Post-translational modifications (PTMs) of the core ATGs regulate their assembly into protein complexes, which execute the sequential steps of autophagosome biogenesis in a timely manner. For example, the ubiquitin-like ATG8 protein undergoes a series of PTMs critical for elongation of the nascent autophagosomal membrane, cargo sequestration and recruitment of other core ATGs (**Fig. 1A** and **B**). The C-terminal peptide of ATG8 is removed by the dedicated protease ATG4, in a process called priming, producing the most abundantly present adduct, ATG8 ΔC. The primed ATG8 (ATG8 ΔC) has its C-terminal glycine (Gly) residue exposed for subsequent modifications^4^. Upon upregulation of autophagy, the exposed Gly of ATG8 ΔC is conjugated with the lipid phosphatidylethanolamine (PE) present on the nascent autophagosomal membranes (**Fig. 1B**) ^4,5^. The ATG8–PE conjugate decorates the inner and outer membranes of the forming autophagosome, playing crucial roles in cargo sequestration, as well as in the formation, closure, and fusion of autophagosomes^6^. At the later stages of vesicle biogenesis, ATG4 associates with the outer membrane of the autophagosome where it cleaves the amide bond between ATG8 and PE, thereby releasing back into the cytoplasmic pool the ATG8 ΔC together with other core ATGs recruited to the membrane by ATG8 (**Fig. 1A** and **B**).

**Figure 1.**
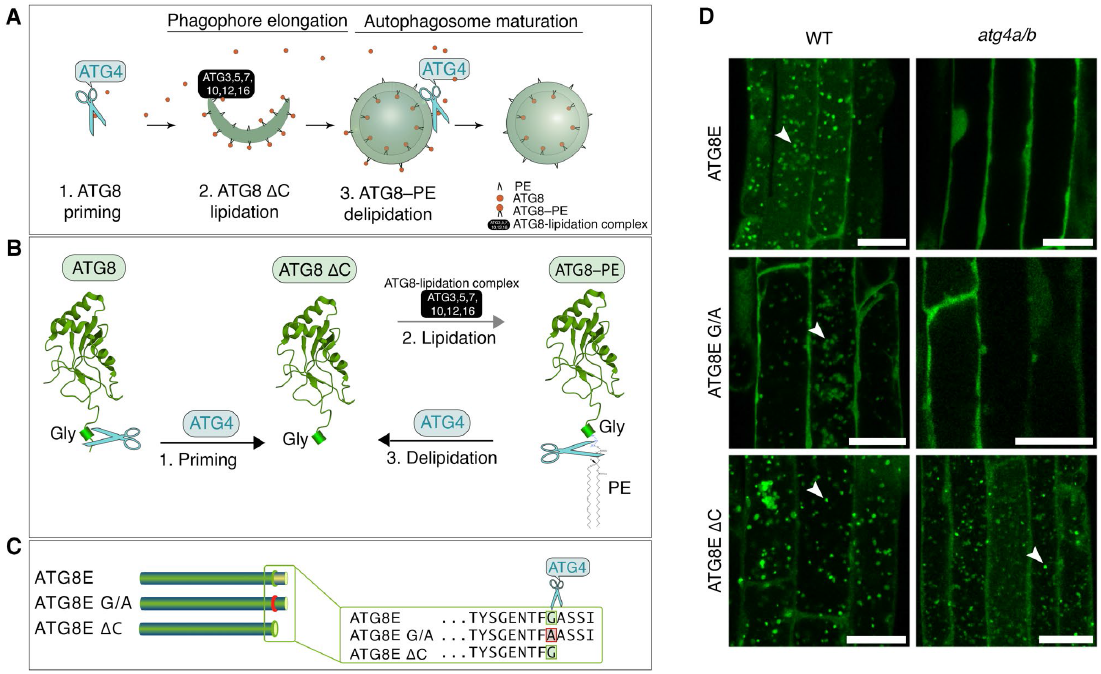
Post-translational modifications of Arabidopsis ATG8 during autophagy. **A**. During autophagosome formation, ATG8 undergoes reversible conjugation with lipids present on the autophagosomal membranes. Lipidation of ATG8 onto phagophore membranes is essential for phagophore elongation. This process begins with the priming of ATG8 through proteolytic processing by the ATG4 protease followed by lipidation aided by other core ATG proteins. Later, lipidated ATG8 is removed from the outer autophagosomal membrane during the maturation of the vesicle. **B**. ATG8 is constitutively primed by ATG4 to remove the C-terminal peptide. The obtained truncated ATG8 (ATG8 ΔC) has the C-terminal Gly residue exposed for the lipidation reaction. Lipidation of ATG8, i.e., its covalent conjugation to the phosphatidylethanolamine (PE) lipid present in the phagophore membranes (formation of ATG8–PE), is executed by ATG8-lipidation complex, best known as two ubiquitin-like conjugation systems, comprising several autophagy-related proteins, including ATG5 and ATG7. Delipidation of ATG8–PE on the outer membrane of autophagosome is performed by the ATG4 protease, which cleaves the amide bond between PE and ATG8, releasing the primed ATG8 ΔC back into the pool of cytoplasmic ATG8 available for autophagosome biogenesis. **C**. Diagram showing C-terminal sequences of the three artificially created variants of the Arabidopsis isoform ATG8E: ATG8E, full-length protein sequence including the C-terminal peptide masking the critical Gly residue; ATG8E G/A, full length protein in which the Gly residue was replaced with Ala; ATG8E ΔC, artificially primed ATG8 protein lacking the C-terminal peptide that was masking the Gly residue. **D**. Detection of three fluorescently labelled ATG8E variants (shown in **C**) in *Arabidopsis thaliana* epidermal root cells of 7-day-old seedlings treated with AZD8055 and concanamycin A. WT, wild-type; *atg4a/b, atg4a-2/b-2* double knockout of ATG4A and ATG4B genes. Arrowheads indicate autophagic bodies inside the vacuoles. The experiment was repeated twice. Scale bars, 20 μm.

This process has two major functions: it facilitates reuse of the released proteins for formation of new autophagosomes, and it modifies the protein shell of the maturing autophagosome^7–9^. Delipidation of ATG8 has been reported to be critical for autophagy in yeast, animals, and plants^6,10,11^. Intriguingly, disruption of this conserved step stalls different stages of autophagosome biogenesis in yeast and mammalian cells. Namely, ATG4-dependent ATG8 delipidation in animal cells is critical for autophagosomal docking and fusion with lysosomes^6,12^. However, abrogation of ATG8 delipidation in yeast impairs an earlier step of autophagosomal membrane elongation^10,13^. This discrepancy indicates fundamental differences in the maturation of yeast and animal autophagosomes and calls for further research necessary for mechanistic comprehension of the variation in the autophagosome biogenesis process among different eukaryotes.

We initiated this study with the aim of elucidating what step of autophagosome biogenesis in plants is affected by the lack of ATG8 delipidation. To our surprise, we discovered that autophagy of the vascular plant *Arabidopsis thaliana* can be carried out to its completion without ATG8 delipidation. Here we provide a thorough verification of the observed phenomenon combining cell biology, biochemistry and reverse genetics, and confirm the physiological relevance of the results by plant phenotyping. Additionally, we show that delipidation remains critical for autophagy of the evolutionarily distant to Arabidopsis unicellular green algae *Chlamydomonas reinhardtii*, that possesses simpler autophagic machinery compared to vascular plants. Lastly, we demonstrate that unlike artificially primed Arabidopsis ATG8E ΔC and ATG8F ΔC isoforms that can fully rescue autophagic flux in the absence of ATG4 activity, the natively primed ATG8I isoform can rescue autophagic activity only partially. These results provide new insights into ATG8 isoform-specific roles in plant autophagosome formation.

## Results

### Expression of ATG8E ΔC restores accumulation of vacuolar puncta in ATG4-deficient Arabidopsis

To identify the specific stage in the process of plant autophagosome formation where ATG8 delipidation plays a crucial role, we produced ATG8-based autophagosomal fluorescent markers in the wild-type (WT) and ATG4-deficient (*atg4a/b*) backgrounds of the vascular plant model organism *Arabidopsis thaliana*. The Arabidopsis genome codes for nine orthologs of ATG8, referred to as isoforms ATG8A–ATG8I^14^. Since the ATG8E isoform has been previously successfully used as a marker for autophagic activity in Arabidopsis^15,16^, we implemented it for this study as well. We generated stable Arabidopsis lines expressing three variants of EosFP-tagged ATG8E: a full-length ATG8, a G/A lipidation-deficient mutant with the critical Gly residue substituted to Ala, and an artificially primed ATG8 ΔC, a truncated protein lacking the C-terminal peptide that would normally mask the critical Gly residue from lipidation (**Fig.1B** and **C**). When expressed in the *atg4a/b* background, ATG8 ΔC should readily form the ATG8–PE adduct anchored in the nascent autophagosomal membranes, however, the latter should not be delipidated in the absence of ATG4 activity.

The seedlings of the established marker lines were simultaneously treated with the autophagy-inducing compound AZD8055 (AZD), and with the compound concanamycin A (ConA) which increases vacuolar pH resulting in deactivation of hydrolases and therefore preservation of the autophagosomes delivered to the plant vacuoles^17^. Expectedly, upon upregulation of autophagy in the WT plants, full-length ATG8E was incorporated into autophagosomes and delivered to the vacuoles, where it was detectable in the form of fluorescent puncta (**Fig. 1D, Fig. S1**). The same marker remained dispersed throughout the cytoplasm in the *atg4a/b* background upon upregulation of autophagy, confirming the critical importance of ATG4-dependent processing for the ATG8’s involvement in autophagosome formation. Lower intensity puncta were observed in the vacuoles of WT cells expressing the G/A mutant of ATG8, indicating that it might be taken up from the cytoplasm as a cargo. Such puncta were not observed in the vacuoles of *atg4a/b* cells, corroborating the lack of autophagic activity in this genetic background. Surprisingly, the artificially primed version, ATG8E ΔC, was observed in the vacuolar puncta of both WT and *atg4a/b*, suggesting that the absence of ATG8 delipidation in the latter case did not impede autophagic activity.

Given the unexpected nature of the observation described above, we found it necessary to confirm that the *atg4/b* mutant background used for this study is indeed ATG4-deficient. For this, we performed a thorough verification of the plants using genotyping, RT-PCR, and qPCR followed by an *in planta* ATG8-cleavage assay (**Fig. S2**). The results of these assays confirmed the presence of the expected T-DNA insertions disrupting *ATG4A* and *ATG4B* genes (**Fig. S2A**), leading to the loss of the corresponding transcripts (**Fig. S2C**) and, subsequently, to the loss of ATG4 proteolytic activity (**Fig. S2E-G**). qPCR analysis revealed a 7-fold increase in the expression of full-length ATG8E in *atg4a/b* (**Fig. S2D**) yet no autophagic bodies were detectable in these plants (**Fig. 1D**). In contrast, a relatively lower expression of the artificially primed ATG8E ΔC (**Fig. S2D)** was sufficient for the formation of the autophagic bodies in the ATG4-deficient background (**Fig. 1D)**.

### Formation of plant autophagosomes and their delivery to the vacuole do not depend on ATG8 delipidation

To verify that the EosFP-positive puncta observed in the vacuoles of *atg4a/b* plants were indeed autophagic bodies labelled with the EosFP–ATG8–PE adduct, we expressed EosFP–ATG8E ΔC in the background lacking ATG8 lipidation activity, i.e., knockout of *ATG5* gene^18^. Transgenic plants expressing the artificially primed version of ATG8E were subjected to AZD/ConA treatment prior to imaging the root and shoot epidermal cells. Upon treatment, the marker localized to vacuolar puncta in the WT and *atg4a/b*, but not in the *atg5* plants, confirming that punctate localization of the marker was dependent on ATG8 lipidation and therefore represented autophagosomal structures (**Fig. 2A**). Quantification of puncta revealed that autophagic bodies accumulated in the vacuoles of *atg4a/b* cells in amounts comparable to WT, indicating reconstituted autophagic activity in the absence of the ATG8-delipidation step (**Fig. 2B**). Furthermore, we conducted a similar analysis on true leaves and confirmed that the observed phenomenon was not restricted to the early developmental stages of Arabidopsis but also occurred in adult plants (**Fig. S3A** and **B**).

**Figure 2.**
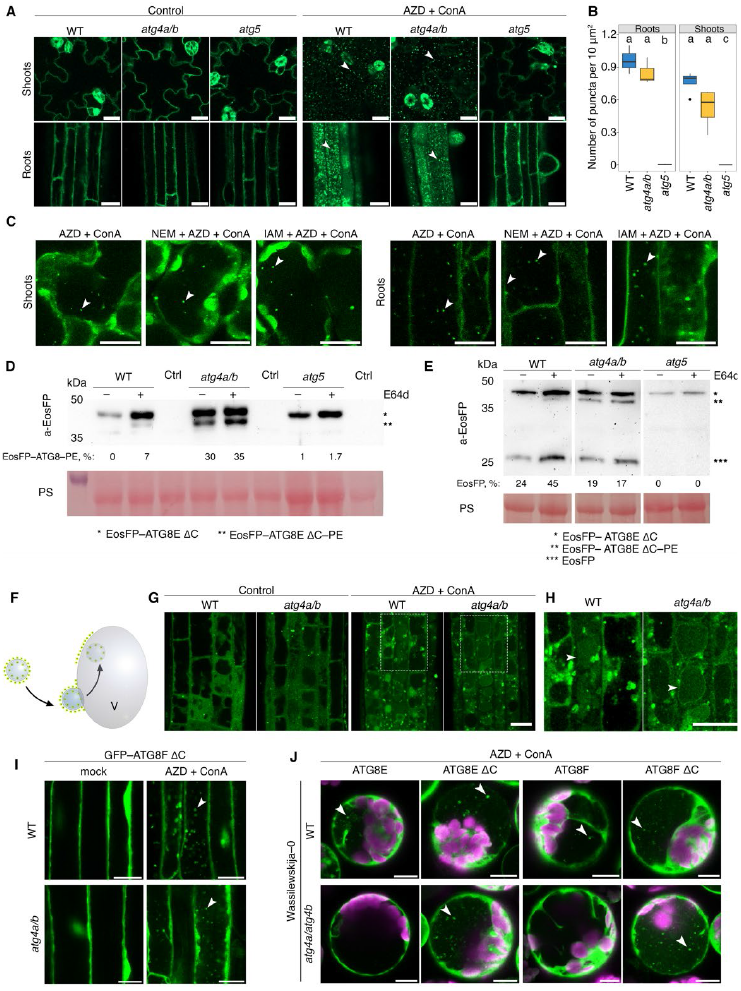
Delipidation of ATG8E and F is dispensable for formation of autophagosomes and their delivery to the vacuole. **(A-H)** demonstrate the impact of expressing the artificially primed ATG8E isoform (ATG8E ΔC) in the *atg4a-2/b-2* knock-out. **(I)** illustrates the similar effect of the artificially primed ATG8F isoform in the *atg4a-2/b-2* knockout. (**J**) depicts the analogous effect of primed ATG8F and ATG8E isoforms in the *atg4a-1/atg4b-1* knockout. **A**. Confocal microscopy images of root and shoot epidermal cells of 7-day-old Arabidopsis seedlings expressing EosFP-ATG8E ΔC in WT, ATG4- or ATG5-deficient backgrounds. Seedlings were incubated under control conditions or subjected to AZD/ConA treatment. Scale bars, 20 μm. **B**. Quantification of puncta per vacuolar area for the AZD/ConA-treated samples illustrated in (**A)**. Tukey’s HSD test was performed for each plant organ individually, *n* = 12 biological replicates (400 technical replicates) for shoots and *n* = 9 biological replicates (210 technical replicates) for roots. Groups that do not differ are annotated by the same letter (α = 0.05). Boxes indicate interquartile range (IQR); median is indicated by a horizontal line; whiskers represent range within 1.5 IQR; outlier is represented by a filled circle. The chart shows representative results of one out of two experiments. **C**. Confocal microscopy images of EosFP–ATG8E ΔC expressed in mesophyll cell of WT Arabidopsis true leaves (left) or roots of 10-day-old seedlings (right) treated with AZD/ConA in the absence or presence of irreversible cysteine protease inhibitors (1mM IAM or 10 mM NEM). Images of leaves were taken at 24 h of treatment, while roots where imaged after 4h of treatment. Scale bars, 20 μm. Time-lapse data confirming that the observed puncta are indeed freely in the vacuoles of leaves is available in **Movie S1**. Quantification of puncta is available in **Fig. S3C**. The inhibitory effect of NEM and IAM treatments on the proteolytic activity of ATG4 was confirmed by Western blot (**Fig. S3D**). **D**. Detection of EosFP in the protein extracts of seedlings treated with AZD/ConA and E64d (illustrated in **Fig. S3C**). Ponceau S staining was used as a loading control. Ctrl, negative control (extract of a WT plant with no transgene). Western blot reveals accumulation of the EosFP–ATG8E ΔC–PE in ATG4-deficient background or in WT upon E64d treatment. Numbers under Western blot lanes represent densitometry results corresponding to integrated density of the lipidated protein band expressed as % of total signal intensity for the corresponding sample. PS, Ponceau S staining used as a loading control. **E**. EosFP cleavage assay performed on the same material used for (**D**) shows no decrease in autophagic activity in the samples accumulating ATG8–PE. Numbers under Western blot lanes represent densitometry results corresponding to integrated density of the free EosFP protein band expressed as % of total signal intensity for the corresponding sample. PS, Ponceau S staining used as a loading control. **F**. Schematic representation of the expected EosFP signal accumulation on the tonoplast upon its fusion with EosFP–ATG8E ΔC–decorated autophagosomes in the absence of ATG8-delipidation activity. **G**. Confocal microscopy images of root epidermal cells expressing EosFP–ATG8E ΔC and showing accumulation of tonoplastic EosFP signal in ATG4-deficeint background upon upregulation of autophagy. Scale bar, 20 μm. **H**. Zoomed-in images of selected areas in (**D)** with adjusted contrast. White arrowheads point at the vacuoles Scale bar, 20 μm. **I**. Epidermal root cells of 7-day-old WT or *atg4a-2/b-2* Arabidopsis seedlings expressing artificially primed version of the ATG8F isoform fused with GFP. Seedlings were imaged under standard conditions (mock) or after 2 h treatment with AZD/ConA. White arrowheads point at autophagic bodies accumulating in the vacuoles upon drug treatment. Scale bar, 15 μm. Quantification of autophagic bodies is available in **Fig. S4B. J**. Confocal images of mesophyll protoplasts isolated from 3-week-old Wassilewskija-0 ecotype WT or *atg4a-1/4b-1* plants and transiently expressing GFP fusions of full length and artificially primed versions of ATG8E and ATG8F isoforms. Protoplasts were subjected to 24 h treatment with AZD/ConA prior to imaging. White arrowheads point at autophagic bodies accumulating in the vacuoles of cells expressing primed versions of ATG8E and ATG8F when treated with drugs to induce autophagy. Scale bar, 10 μm. The complete data set including untreated control is available in **Fig. S6**.

Next, we corroborated the results obtained *via* genetic ablation of ATG4 by using chemical inhibitors of ATG4 activity. For this, 10-day-old seedlings and one-month-old plants expressing EosFP-ATG8E ΔC in the WT background were treated with AZD and ConA supplemented with cysteine protease inhibitors iodoacetamide (IAM) or N-ethylmaleimide (NEM), chemical compounds previously shown to irreversibly inhibit ATG4 activity^19–21^. Neither IAM nor NEM treatment prevented accumulation of autophagic bodies in the vacuoles of roots or shoots (**Fig. 2C; Fig. S3C and D, Movie S1**), although some decrease in the number of puncta was observed in highly stressed cells. Due to the strong cytotoxicity of these chemical compounds, we decided to adhere to the use of genetic mutants for further *in planta* experiments. Additionally, we tested another inhibitor of cysteine proteases, E64d which was suggested to not impact ATG4 activity but potentially stabilize autophagic bodies in the lytic compartment^22^. Accordingly, we did not observe a significant difference between WT and *atg4a/b* mock- or E64d-treated plants expressing EosFP–ATG8E ΔC (**Fig. S3E** and **F**). Western blot detection of EosFP–ATG8E ΔC in the plants subjected to AZD/ConA treatment with or without E64d revealed accumulation of the ATG8–PE adduct in the ATG4-deficient background independently of treatment. However, treatment with E64d rendered ATG8–PE detectable in the WT protein extracts, most probably due to the better stability of the adduct under these conditions (**Fig. 2D**). The ATG8–PE band was not detectable in the protein extracts of *atg5*, consistent with the lack of ATG8-lipidating activity in this mutant (**Fig. 2D**). Finally, we validated that the lower band identified in Western blots indeed represents the ATG8–PE adduct by conducting an *in vitro* ATG8 delipidation assay (**Fig. S3G** and **H**).

A study carried out in yeast revealed an APEAR motif (ATG8–PE association region) responsible for ATG4 recruitment to the lipidated form of ATG8 during autophagosome maturation^9^. Intriguingly, the APEAR motif is conserved in the ATG4B but not the ATG4A Arabidopsis ortholog^9^, indicating potential functional diversification of these orthologs in ATG8 processing. Importantly, individual knockouts of *ATG4* in Arabidopsis do not display a discernible autophagy-deficient phenotype^8,23^ and accumulate autophagic bodies in WT-like quantities upon AZD/ConA treatment (**Fig. S3I** and **J**). If the two ATG4 enzymes indeed differ in their contribution to ATG8 delipidation, the lack of distinct phenotypes in the corresponding knockouts could provide further evidence for ATG8 delipidation not being critical for autophagy. To verify this hypothesis, we tested if delipidation efficacy is reduced in the single knockout for *ATG4B* bearing APEAR motif but not in the single *ATG4A* knockout when compared to WT. For this, we crossed *atg4a/b* and WT plants expressing EosFP–ATG8E ΔC and performed Western blot detection of the tagged ATG8 in the young (2-week-old) and older (6-week-old) plants. To our surprise, delipidation was significantly reduced in both individual *ATG4* knockouts (**Fig. S3K** and **L**). Despite an at least 4-fold decrease in ATG8-delipidating activity in the single ATG4 knockouts compared to WT (**Fig. S3K** and **L**) no difference in the number of autophagic bodies was observed in these backgrounds upon induction of autophagy (**Fig. S3I** and **J**). These results are in agreement with the other observations pointing towards ATG8 delipidation being dispensable for plant autophagy. They also demonstrate that while the proteolytic activity required for C-terminal processing of ATG8 can be efficiently executed by a single ATG4 isoform (**Fig. S2E** and **F**), the delipidating activity of individual Arabidopsis ATG4s is significantly less efficient and requires expression of both genes.

To corroborate the reconstitution of autophagic activity in the absence of ATG8 delipidation, we deployed a modified version of the GFP cleavage assay^22^, involving detection of EosFP– ATG8 processing upon induction of autophagy with AZD/ConA/E64d treatment. The assay demonstrated efficient processing of the fusion protein and accumulation of free EosFP in the plants expressing EosFP–ATG8E ΔC in the WT and *atg4a/b*, but not in the *atg5* background (**Fig. 2E**), thereby corroborating restored autophagic activity in the absence of ATG8 delipidation.

We hypothesized that lack of ATG8-delipidating activity would result in a large number of EosFP–ATG8E ΔC-decorated autophagosomes fusing with the vacuolar membrane, which should eventually yield fluorescent labeling of the tonoplast (**Fig. 2F**). Indeed, induction of autophagy caused accumulation of tonoplastic EosFP signal in the *atg4a/b*, but not in the WT cells (**Fig. 2G and H)**.

To confirm that the above-described observations were not specific to ATG8E, we generated transgenic lines expressing an artificially truncated version of another highly expressed Arabidopsis ATG8 isoform, ATG8F ΔC, in the WT and *atg4a/b* backgrounds. Indeed, treatment of these plants with AZD/ConA triggered a similar accumulation of autophagic bodies in both backgrounds, consistently observed in the roots of seedlings and true leaves of 4-week-old plants (**Fig. 2I, Fig. S4A** and **B**). qPCR analysis of the established lines showed a slightly lower expression level of the transgene in the *atg4a/b vs*. WT background (**Fig. S4C**), which still yielded a comparable quantity of autophagic bodies (**Fig. S4B**) and GFP-cleavage efficacy (**Fig. S4D**) in both backgrounds. In line with our findings from ATG8E ΔC-expressing plants, increased levels of ATG8F–PE were observed in the *atg4a/b* background (**Fig. S4E**), illustrating defect in ATG8 delipidation.

Finally, we tested the reproducibility of the observed effect in an alternative ATG4-deficient background, specifically in the *atg4a-1/atg4b-1* knockout, which was utilized in the original article investigating the role of ATG8 delipidation in Arabidopsis autophagy^8^. For this, we extracted mesophyll protoplasts from the leaves of *atg4a-1/atg4b-1* mutants and the WT plants of the corresponding Wassilewskija-0 ecotype. The protoplasts were transformed to express GFP-tagged full-length or artificially primed ATG8E or ATG8F isoforms, treated with AZD/ConA and imaged using confocal microscopy (**Fig. 2J** and **Fig. S5**). We observed that the expression of artificially primed ATG8 isoforms restored the accumulation of autophagic bodies also in this ATG4-deficient background.

In summary, we found that while, as expected, the ATG4-deficient plants expressing artificially primed ATG8 isoforms accumulate ATG8–PE adduct, these plants, unexpectedly, do not exhibit any defects in the formation or vacuolar delivery of autophagosomes.

### ATG8 delipidation-independent autophagy restores wild-type like stress tolerance in the ATG4-deficient plants

While we confirmed the normal formation of plant autophagosomes in the absence of ATG8 delipidation, it still remained a question of whether such autophagosomes could carry out their intended function. To test this, we examined the stress tolerance of *atg4a/b* plants expressing EosFP–ATG8E ΔC under macronutrient-deprived conditions which would rely on autophagic activity^1,3^ (**Fig. 3A-F**).

**Figure 3.**
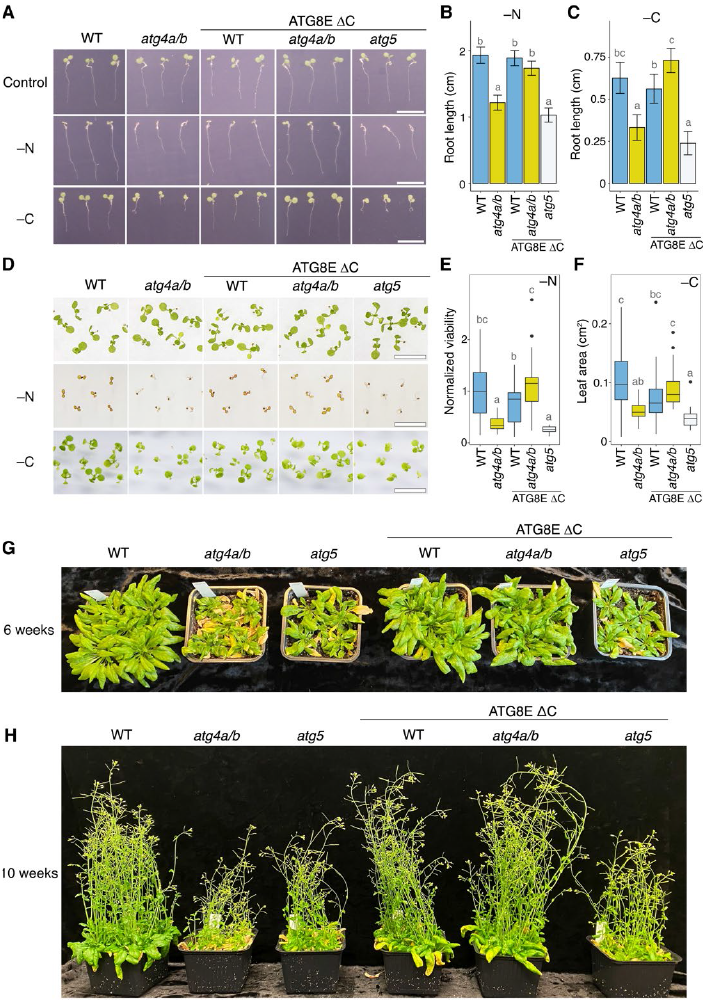
Expression of ATG8E ΔC alleviates autophagy-deficient phenotypes of *atg4a/b*. **A**. Examples of Arabidopsis seedlings phenotyped using SPIRO under control, nitrogen-depleted (–N), and carbon-depleted (–C) conditions. Normal root growth under nutrient-depleted conditions is restored in *atg4a/b* seedlings expressing ATG8E ΔC. In total, 997 biological replicates were phenotyped under –N and –C conditions in seven independent experiments. Scale bars, 1 cm. **B**. Quantification of seedling root length on the 4^th^ day after germination when grown under –N conditions, as illustrated in panel **A** (middle row). The chart represents combined data from two independent experiments, *n* = 364. Root growth was analyzed by fitting a mixed-effect second-order polynomial model to the time-resolved root length data obtained by the SPIRO root growth assay. Letters signify all-pairwise comparisons, where treatments sharing the same letter do not differ at α = 0.05. Error bars indicate 95% confidence interval. **C**. Quantification of seedling root growth during four days of recovery following four days of carbon depletion, as illustrated in panel **A** (bottom row). The chart represents combined data from two independent experiments, *n* = 283. Root length was analyzed and visualized as in (**B**). **D**. Examples of Arabidopsis seedling shoot phenotypes under control, nitrogen-depleted (–N), and carbon-depleted (–C) conditions. Scale bars, 1 cm. Photos of complete plates are shown in **Fig. S7. E**. Viability of Arabidopsis seedlings grown on –N medium illustrated in (**D**, middle row) was estimated by assessing shoot chlorosis. Phenotyping was performed in two independent experiments. The chart summarizes data from a single experiment, *n* = 157. The upper and lower edges of boxes indicate upper and lower quartiles of the data, the middle line indicates the median, and the whiskers extend to data points within 1.5 * interquartile range (IQR). Outliers outside 1.5 * IQR are represented by filled circles. Tukey’s HSD test, shared letters indicate no difference at α = 0.05. **F**. Efficacy of recovery after carbon-depletion illustrated in (**D**, bottom row) was estimated by measuring the leaf area for each seedling at the end of a one-week-long recovery after 7 days of carbon depletion. The experiment was performed twice. Chart shows representative data from one experiment, *n* = 160. Results were analyzed and visualized as in (**E**). **G**. Rosette phenotype of 6-week-old plants grown under long day conditions show a typical attribute for autophagy-deficiency: an early onset of senescence in the *atg4a/b* and *atg5* backgrounds, and lack thereof in the WT and *atg4a/b* plants expressing ATG8E ΔC protein. **H**. Inflorescence phenotype imaged one month later for the same plants as shown in **G**.

We performed uninterrupted time-lapse imaging of seedlings under nitrogen-deprived (–N) and carbon-deprived (–C) conditions. For this, seeds of WT, *atg4a/b* and *atg5* plants expressing EosFP–ATG8E ΔC were plated on standard 0.5xMS growth medium, and on growth media depleted of either nitrogen or sucrose. Plates were photographed every hour for 7 days using a SPIRO imaging platform^24^. For –C starvation induction, imaging was performed in the dark growth chamber using SPIROs green LED for illumination during image acquisition, to prevent photosynthetic carbon assimilation. We observed stagnated root growth in autophagy-deficient plants (*atg4a/b* and *atg5*) under both – N and –C conditions (illustrated in **Fig. 3A-C**; time-lapse movies showing complete plates are available in **Movies S2** and **S3**). Expression of EosFP–ATG8E ΔC in the WT background did not have a discernible effect on root growth, however, it reconstituted WT-like root growth in the *atg4a/b* background, revealing functional autophagic activity in these plants. No such effect was observed in the *atg5* plants expressing EosFP–ATG8E ΔC. These results were corroborated by our observations that the expression of another artificially primed ATG8 isoform, ATG8F, reinstated the tolerance of the *atg4a/b* mutants to nitrogen depletion (**Fig. S4F** and **G**). Similar to root phenotyping, assessment of seedling shoot viability and leaf growth under the same stress conditions revealed restored stress-tolerance in the *atg4a/b* upon expression of ATG8E ΔC (**Fig. 3D-F**). Uncropped images of plates with plants used in these experiments are shown in **Fig. S6**.

Finally, we phenotyped the transgenic plants expressing artificially primed ATG8E and ATG8F isoforms at the later developmental stages. Plants were grown in soil under long day conditions to detect the early senescence phenotype typical for autophagy-deficient Arabidopsis plants^8^. Consistent with the above-described phenotyping assays, we could observe alleviation of the early senescence phenotype in the *atg4a/b* plants expressing either of the two truncated ATG8 isoforms (**Fig. 3G** and **H, Fig. S4H** and **I**).

In summary, the phenotypic analyses of stress-tolerance of the photosynthetic organs and roots, carried throughout the life span of Arabidopsis, indicate that autophagic structures formed in the absence of ATG8 delipidation correspond to fully functional autophagosomes.

### Arabidopsis ATG8 isoforms differ in their capacity to carry out autophagic flux

Two out of nine Arabidopsis ATG8 isoforms, ATG8H and ATG8I, lack the C-terminal peptide and therefore do not require processing by ATG4 for their lipidation. It is conceivable that these two isoforms can be readily lipidated and used for biogenesis of autophagosomes in the *atg4a/b* mutant (**Fig. 1A** and **B, Fig. 4A**). Reflecting upon our discovery that autophagy can be re-established in the ATG4-deficient background through expression of the artificially primed ATG8E or ATG8F, we wondered why the natively primed ATG8 isoforms do not similarly sustain autophagosome formation in *atg4a/b* plants. We considered two hypotheses: (i) the sum amount of ATG8H and ATG8I (which would correspond to the total amount of ATG8 available for lipidation in *atg4a/b*) is much lower than the sum amount of all isoforms ATG8A–ATG8I available for lipidation in the WT background, and might not be sufficient for normal autophagosome biogenesis; (ii) ATG8H and ATG8I isoforms play specific roles in plant autophagy that do not allow efficient autophagosome biogenesis in the absence of other isoforms, e.g., sequestration of specific cargo. Notably, both ATG8H and ATG8I show higher protein sequence similarity with animal ATG8 orthologs compared to ATG8A–G (**Fig. S7A** and **B**)^14^, suggesting a potential functional diversification of these two groups of the Arabidopsis ATG8 isoforms.

**Figure 4.**
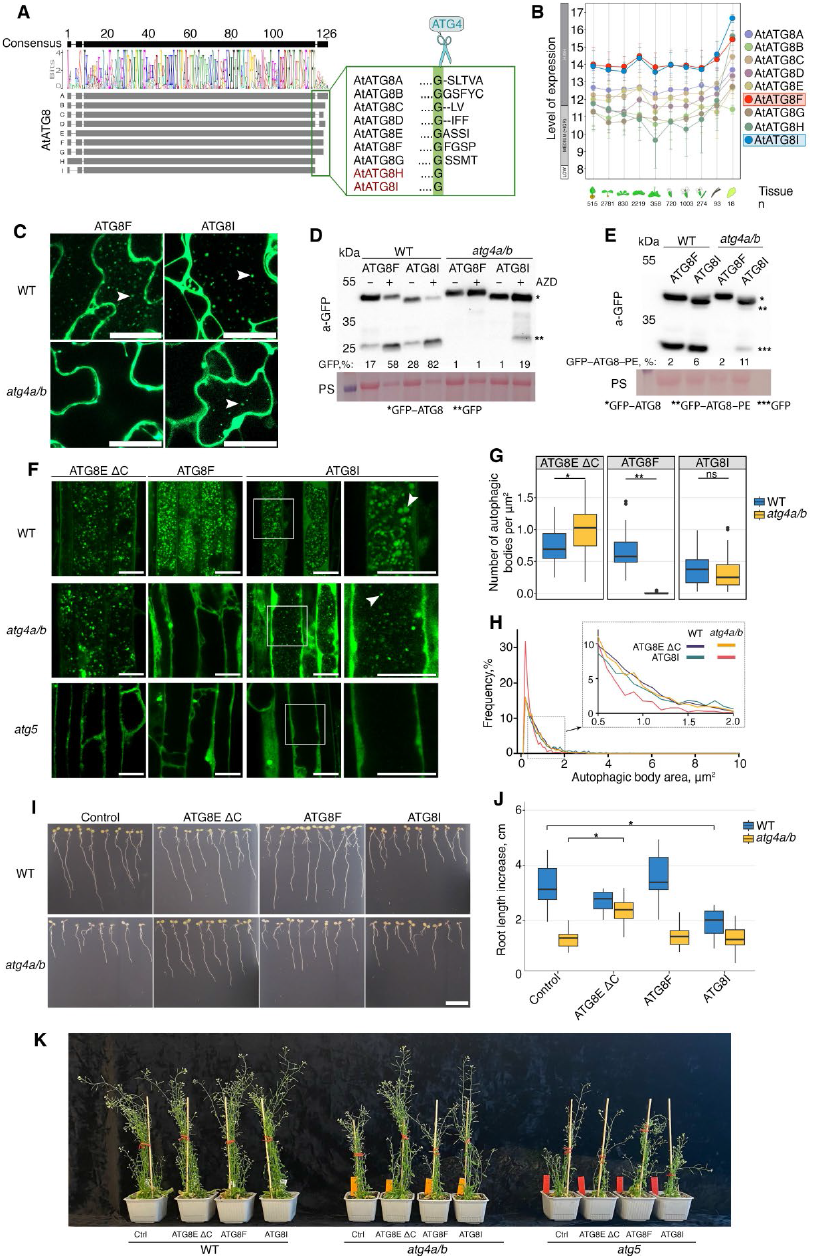
Arabidopsis ATG8 isoforms differ in their capacity to carry out autophagic flux. **A**. Schematic representation of ClustalW protein alignment of the nine Arabidopsis ATG8 isoforms. The inset shows ATG8 C-terminal sequences containing the Gly residue which is critical for lipidation. Notably, ATG8H and ATG8I do not possess the C-terminal peptide and are therefore natively primed ATG8 isoforms having the critical Gly readily exposed. **B**.GENEVESTIGATOR data on organ-specific expression of all nine ATG8 isoforms of Arabidopsis. ATG8F (red rectangle) and ATG8I (blue rectangle) isoforms are expressed at the highest level in all tissues. **C**.Confocal microscopy images showing accumulation of autophagic bodies in the vacuoles of leaf epidermal cells stably expressing GFP-tagged ATG8F or ATG8I in WT or *atg4a/b*. Leaves were infiltrated with AZD/ConA 24 h prior to imaging. Two independent lines were checked for each genotype. White arrowheads point at autophagic bodies. Scale bars, 25 μm. **D**.GFP cleavage assay performed on the transgenic lines illustrated in **C**, showing restored accumulation of free GFP in the *atg4a/b* plants expressing GFP–ATG8I. Autophagy was induced by infiltrating leaves with AZD 24 h prior to sampling. Densitometry results are shown as numbers under corresponding lanes and represent integrated density for the free GFP protein band expressed as % of total signal detected in the corresponding sample. PS, Ponceau S was used as a loading control. **E**. Western blot detection of GFP in AZD-treated samples shown in **D** separated on the gel containing 6 M urea to discern lipidated from non-lipidated forms of ATG8 confirms accumulation of the lipidated ATG8I. The panel shows results representative for two independent experiments. Densitometry results are shown as numbers under corresponding lanes and represent integrated density for the lipidated ATG8 protein band expressed as % of total signal detected in the corresponding sample. PS, Ponceau S was used as a loading control. **F**. Confocal microscopy images showing accumulation of autophagic bodies in root epidermal cells of seedlings expressing three isoforms of Arabidopsis ATG8 (ATG8E ΔC, ATG8F and ATG8I) in the WT, ATG4- or ATG5-deficient backgrounds. Seven-day-old Arabidopsis seedlings were subjected to AZD/ConA treatment for 24 h prior to imaging. Scale bars, 20 μm. White arrowheads point at autophagic bodies in the vacuolar lumen. **G**. Quantification of autophagic bodies in vacuoles of cells illustrated in **F**. Two independent lines were checked for each of the ATG8I- and ATG8F-expresing background. White Welch’s t-test, *n* = 337. *, *p* < 0.05; **, *p* < 0.01; ns, not significant. Boxes indicate interquartile range (IQR); median is indicated by a horizontal line; whiskers represent range within 1.5 IQR; outliers are represented by filled circles. **H**. Size distribution of autophagic bodies in the samples illustrated in **F** and quantified in **G**. The inset shows a magnified portion of the chart visualizing the higher frequency of smaller autophagic bodies present in the vacuoles of *atg4a/b* plants expressing ATG8I. **I**. Root growth phenotype of 12-day-old seedlings incubated for 7 days on nitrogen depleted medium. In contrast to the artificially primed ATG8E ΔC, expression of the natively truncated ATG8I isoform is not sufficient to alleviate the autoph-agy-deficient phenotype of *atg4a/b*. Two independent lines were checked for each of the ATG8I- and ATG8F-expresing background. Seedlings were grown for 4 days on standard 0.5xMS medium, checked for GFP signal, then transferred onto –N medium plates to be imaged for 7 days. Scale bar, 1 cm. **J**. Quantification of the increase in root length of the seedlings after 7 days growth on the –N medium (illustrated in **I)**. The experiment was performed three times. Boxes indicate interquartile range (IQR); median is indicated by a horizontal line; whiskers represent range within 1.5 IQR. The chart shows representative data from one experiment. Dunnett’s tests were performed for WT and *atg4a/b* separately, comparing each genotype against the control sample, *n* =128. *, *p* < 0.0001. **K**. Phenotypes of two-month-old plants grown in soil under long day conditions. Photos show plants with representative phenotypes, the experiment was performed twice using four biological replicates for each genotype. Four independent lines were checked for each of the ATG8I- and ATG8F-expresing background. Ctrl, a not transformed plant of the corresponding genotype; ATG8E ΔC, ATG8F and ATG8I, overexpression of the fluorescent fusion of the corresponding ATG8 isoform.

To test our hypotheses, we selected ATG8F and ATG8I as the two strongest expressed ATG8 isoforms, one with and one without C-terminal peptide, respectively (**Fig. 4B**). We generated fluorescently labelled fusions of these isoforms and expressed them transiently from a strong native Arabidopsis promoter (*APA1*^25^) in WT, *atg4a/b*, and *atg7* cells, using the latter as a negative control lacking ATG8 lipidation activity. Upon induction of autophagy, we could observe the expected accumulation of autophagic-body-like structures in the vacuoles of WT cells expressing either of the ATG8 isoforms, and lack thereof in the vacuoles of *atg7* mutants (**Fig. S7C**). Intriguingly, overexpression of ATG8I in the *atg4a/b* background seemed to have reconstituted accumulation of puncta in this background, thus supporting our first hypothesis that the amount of ATG8 available for lipidation might be the limiting factor in *atg4a/b* autophagy. As expected, ATG8F overexpression in the same cells did not reconstitute autophagosome biogenesis.

To enable an in-depth investigation of this finding using biochemistry and phenotyping assays, we generated stable transgenic lines overexpressing GFP-fusions of the same two ATG8 isoforms under the control of a strong viral promoter (2×35S), and verified expression levels of the transgenes in the new lines using qPCR (**Fig. S4C and S8D**). Detection of autophagic bodies in the true leaves of these plants confirmed the results obtained in the transient expression system: indeed, overexpression of ATG8I was sufficient to reconstitute accumulation of autophagic bodies in the vacuoles of *atg4a/b* upon AZD/ConA treatment (**Fig. 4C**). The GFP-cleavage assay performed on these transgenic lines further confirmed autophagic flux present in the *atg4a/b* overexpressing the natively truncated ATG8I isoform (**Fig. 4D**).

Intriguingly, detection of the lipidated and non-lipidated adducts of ATG8 in the protein extracts from plants overexpressing ATG8F and ATG8I isoforms revealed accumulation of ATG8I–PE not only in the *atg4a/b* but also in the WT background (**Fig. 4E**), suggesting a decreased affinity of ATG4 towards this isoform.

However, although GFP-cleavage assay revealed the occurrence of autophagic activity in the ATG4-deficient plants overexpressing ATG8I, the decreased cleavage of GFP–ATG8I in the *atg4a/b* background in comparison to the WT (**Fig. 4D** and **E**) suggested that our second hypothesis might also be correct, and that ATG8I cannot fully reconstitute normal autophagosome formation. To verify this further, we compared the number of puncta formed in the Arabidopsis root cells overexpressing ATG8E ΔC, ATG8F, or ATG8I upon upregulation of autophagy. For this, Arabidopsis seedlings over-expressing either of the ATG8 isoforms in the WT, *atg4a/b*, or *atg5* backgrounds were subjected to AZD/ConA treatment for 24 h followed by confocal microscopy (**Fig. 4F**). Quantification of the puncta on the obtained micrographs revealed no significant difference between puncta density in the WT and *atg4a/b* overexpressing ATG8I (**Fig. 4G**), indicating normal rate of autophagosome formation and their delivery to the vacuole. However, it did not escape our notice that the size of the GFP– ATG8I-positive puncta differed in the WT and *atg4a/b* cells. Quantification of the puncta size distribution showed that indeed, autophagic bodies formed in the *atg4a/b* cells overexpressing ATG8I were significantly smaller in comparison to autophagic bodies formed in the vacuoles of WT cells overexpressing the same isoform (**Fig. 4H**), indicating potential abnormalities in the elongation of autophagosomes in the presence of ATG8I alone.

Next, we tested if the decreased size of autophagic bodies in the ATG8I-overexpressing *atg4a/b* plants impacted the efficacy of autophagic activity. We subjected seedlings to nitrogen starvation and measured elongation efficacy of their roots using SPIRO time-lapse images (**Fig. 4I** and **J**). Notably, the overexpression of the natively primed ATG8I did not rescue the stagnated root growth phenotype of *atg4a/b*, indicating that smaller autophagosomes observed in this background were not sufficient to cope with the scarcity of exogenous nitrogen. Surprisingly, under the same conditions, we also observed decreased root elongation in the WT plants overexpressing ATG8I (**Fig. 4I** and **J**).

Considering that the puncta quantification experiments (**Fig. 4F** and **G**) were performed using AZD as a trigger for autophagy, while root phenotyping was done under –N conditions (**Fig. 4I** and **J**), we decided to test if the overexpression of ATG8I has a different effect on WT depending on the stress conditions. For this, we compared density and size of autophagic bodies upon AZD, –N and –C treatments (**Fig. S8**) in the lines overexpressing either ATG8E ΔC, which caused no discernible phenotype in the WT in the previous experiments, or ATG8I which caused decreased tolerance to nitrogen depletion in WT. In agreement with our previous observations, ATG8E ΔC overexpression in the WT and *atg4a/b* backgrounds yielded similar density and size of autophagic bodies under all three stress conditions, while overexpression of ATG8I resulted in smaller autophagic bodies (**Fig. S8A-C**). The latter phenotype was especially prominent under –N conditions (for WT: 0.51 μm^2^ under –N vs 0.76 μm^2^ under –C, or 0.69 μm^2^ under AZD, Tukey’s HSD test *p*-value <0.001; **Fig. S8A-C**), thereby explaining the observed stunted root growth on nitrogen-depleted medium. The autophagic flux rate under the three stress conditions was additionally confirmed using GFP-cleavage assay (**Fig. S8D**). Since ATG8I overexpression reduced the size of autophagic bodies in the WT cells under all three types of stress conditions, it is tempting to speculate that an excessive amount of this isoform could cause premature closure of phagophores.

Interestingly, when plants were grown under favorable long day conditions in soil, WT overexpressing ATG8I showed no discernible phenotype, while *atg4a/b* plants overexpressing the same isoform grew more fit than autophagy-deficient knockouts (**Fig. 4K**). These observations suggest that ATG8I overexpression can rescue autophagy of the *atg4a/b* knockout to a level that is sufficient for WT-like plant fitness under favorable conditions, and that toxic effects of ATG8I overexpression in the WT is specific for certain types of stresses, including –N.

In summary, we demonstrated that the autophagy-deficient phenotype of *atg4a/b* is caused by the insufficient amount of the natively truncated ATG8 isoforms available for lipidation during autophagosome formation. We showed that overexpression of the natively truncated ATG8I is sufficient to reconstitute normal rate of autophagosome formation and their delivery to the vacuole but yields autophagosomes almost twice smaller than those formed when artificially truncated ATG8E isoform is expressed in the same background. Furthermore, overexpression of the natively truncated ATG8I isoform undermines the tolerance of WT plants to nitrogen deprivation, possibly by shifting autophagy selectivity to the one unfavorable under such conditions.

### Ultrastructural analysis confirms that vacuolar puncta observed in ATG4-deficient plants expressing primed ATG8 isoforms indeed represent autophagic bodies

To further verify the nature of the vacuolar puncta accumulating in the Arabidopsis transgenic lines upon induction of autophagy, we performed transmission electron microscopy (TEM). For this, seedlings of the transgenic lines expressing the artificially primed ATG8E ΔC, the natively primed ATG8I, and full-length ATG8F isoforms in the WT and *atg4a/b* backgrounds were subjected to AZD/ConA treatment followed first by confocal laser scanning microscopy (CLSM) and later by TEM (**Fig. 5**). Predictably, for the *atg4a/b* plants, CLSM revealed the accumulation of the fluorescent puncta only in the vacuoles of lines expressing primed versions of ATG8s, whereas vacuolar puncta could be detected for all lines in the WT background. TEM showed that the vacuoles of ATG8E ΔC-expressing lines accumulate numerous vesicles filled with varied cargo, including portions of cytoplasm and organelles. No such vesicles were detectable in the *atg4a/b* cells expressing full-length ATG8F. Notably, vacuolar vesicles observed in the *atg4a/b* cells expressing natively primed ATG8I were noticeably smaller than those observed in the vacuoles of ATG8E ΔC-expressing cells (**Fig. 5**). These findings further support the quantitative data obtained through high-throughput CLSM-based assessment of autophagic body size (**Fig. S8A-C)**.

**Figure 5.**
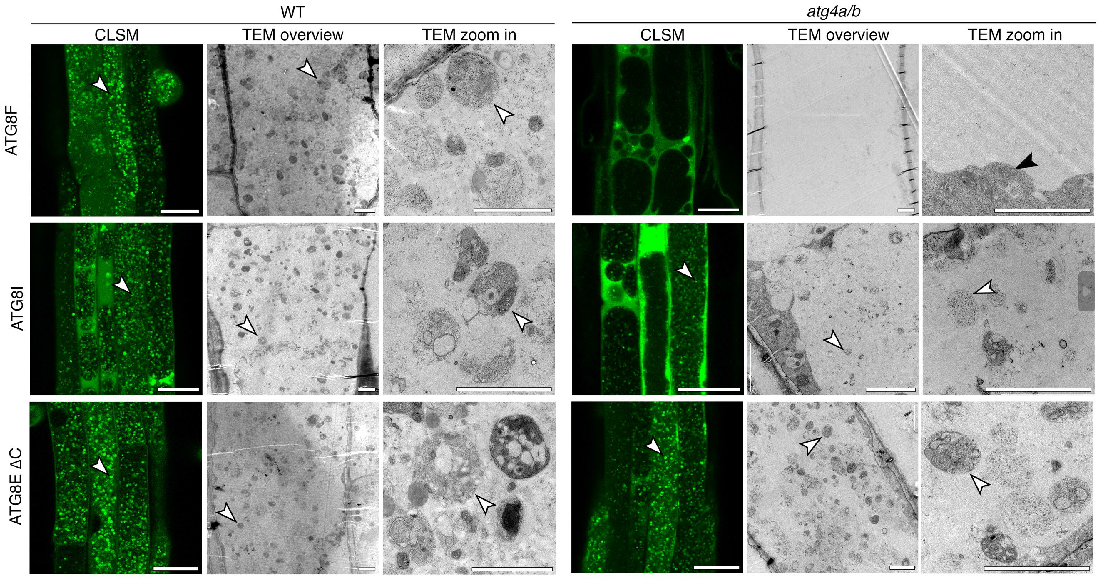
Ultrastructure of GFP-positive puncta accumulating in the vacuoles upon expression of full-length and primed GFP-tagged ATG8 isoforms. Seven-day-old Arabidopsis seedlings expressing GFP-tagged full-length (ATG8F), natively primed (ATG8I) or artificially primed (ATG8E ΔC) ATG8 variants in WT or in *atg4a/b* backgrounds were subjected to AZD/ConA treatment for 2h prior to confocal imaging (CLSM). After CLSM, seedlings were fixed and processed for transmission electron microscopy (TEM). Ultrastructural analysis of the GFP-positive puncta detected with CLSM revealed numerous autophagic bodies with various cargo (cytoplasm, mitochondria, ER). No autophagic bodies were detected in the vacuoles devoid of GFP-positive puncta (*atg4a/b* plants expressing GFP–ATG8F). In agreement with our observations made using CLSM (**Fig. 4H**), autophagic bodies detectable in the *atg4a/b* plants expressing natively primed GFP–ATG8I isoform were visibly smaller than in the *atg4a/b* plants expressing artificially primed GFP–ATG8E ΔC or in WT expressing any of the checked ATG8 isoforms. White arrowheads point at autophagic bodies containing organelles and portions of cytoplasm. Black arrowhead points at mitochondria in the cytoplasm. Scale bar for CLSM images, 25 μm. Scale bars for all TEM micrographs, 2 μm.

In summary, ultrastructural analysis of the vacuolar puncta observed in the transgenic lines confirmed that they represent autophagic bodies.

### ATG8 delipidation is critical for autophagy in a green microalga Chlamydomonas

The current phylogeny categorizes green plants (*Viridiplantae*) as a monophyletic taxon consisting of two evolutionary lineages that diverged over a billion years ago: the Chlorophyta (comprising some of the green algae) and Streptophyta (including both land plants and remaining green alga)^26,27^. To investigate whether independence of autophagy on ATG8 delipidation is inherent for green plants we conducted corresponding experiments on the unicellular green alga, *Chlamydomonas reinhardtii*. Most interestingly, in depth studies of these algae revealed First, we generated *cratg4* and *cratg5* autophagy-deficient mutants in the commonly used that they possess numerous animal-like functions^28,29^. Chlamydomonas genome encodes single genes for the core ATGs, making it a much simpler model for autophagy research compared to Arabidopsis. Intriguingly, the sole ATG8 protein of Chlamydomonas retained its C-terminal peptide, which necessitates the ATG4-mediated priming prior to ATG8 lipidation (**Fig. 1A** and **B**). Therefore, we generated a truncated version of *Chlamydomonas reinhardtii* ATG8, CrATG8 ΔC, which had the critical C-terminal Gly exposed and could be lipidated in the absence of ATG4 activity but would depend on it for the delipidation.

UVM4 background (**Fig. S9**) and complemented them with mCherry-tagged full-length CrATG8 or CrATG8 ΔC. Immunoblot analysis showed a good expression level of both transgenes in the mutant strains (**Fig. 6A**). A lack of autophagy elicits an early senescence phenotype, especially under nutrient-scarce conditions^30,31^. Therefore, we assessed the functionality of the autophagic pathway in the established strains by assessing their rate of senescence. We cultivated the strains in the standard TAP growth medium for 40 days and monitored the degree of their chlorosis while they were gradually depleting nutrients from the medium (**Fig. 6B**). Expectedly, *cratg5* mutants complemented with full-length or truncated ATG8 but still deficient in ATG8 lipidation exhibited chlorosis associated with the early senescence phenotype, while WT cells remained dark green under the same conditions (**Fig. 6B**). Notably, the early onser of chlorosis in the *cratg4* mutant was not rescued by the expression of the truncated ATG8 version (**Fig. 6B**). Taken together, our data suggests that, unlike in the vascular plant, Arabidopsis, but similar to yeast and mammals, the delipidation of ATG8 plays a crucial role in the autophagy of Chlamydomonas.

**Figure 6.**
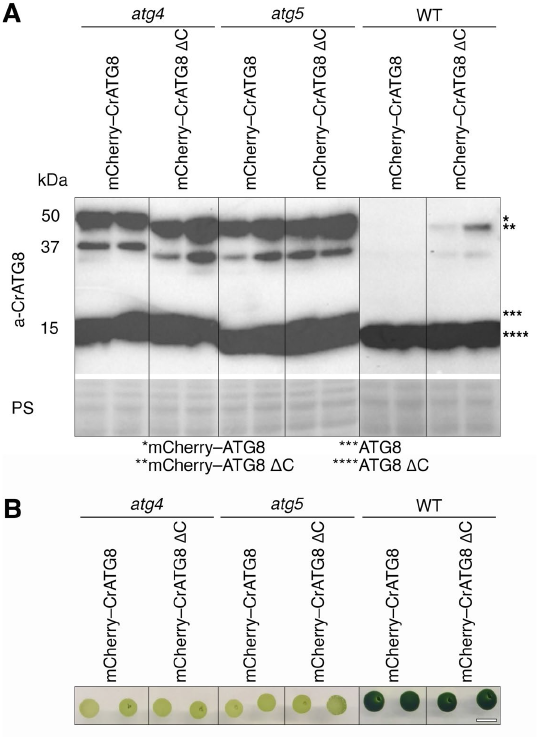
Delipidation of ATG8 is critical for autophagy in *Chlamydomonas reinhardtii*. **A**. Western blot detection of endogenous ATG8 and mCherry–CrATG8 transgene expression in the WT, *cratg4*, and *cratg5* backgrounds. Ponceau S staining was used as a loading control. The cells were grown in the liquid TAP medium under standard conditions and harvested at a density of 1-2×10^7^ cells ml^-1^. **B**. Senescence phenotype of the transgenic *Chlamydomonas* lines grown on the TAP medium for 40 days. Light color of colonies indicates nutrient deficiency sensitivity caused by dysfunctional autophagy. Scale bar, 1 cm.

## Discussion

In this study we demonstrate that ATG8 delipidation is dispensable for autophagy in the vascular plant *Arabidopsis thaliana*, which belongs to the Streptophyta lineage. However, it is of crucial importance for autophagy in the unicellular green alga *Chlamydomonas reinhardtii*, which is part of the Chlorophyta lineage and has preserved a number of animal-like fucntions^28,29^. The specific evolutionary adaptations that rendered delipidation dispensable have yet to be uncovered.

Since ATG8 delipidation was previously reported to be crucial for autophagy in plants^8^, as well as in yeast and animal cells^6,10,11^, we considered it imperative to conduct a thorough verification of our surprising observation regarding the reconstituted autophagosome formation in the ATG4-deficient Arabidopsis. We employed a combination of mutagenesis, cell biology, biochemistry, and phenotyping assays to establish a robust foundation for future investigations aimed to elucidate the molecular mechanisms governing plant-specific maturation of autophagosomes possible in the absence of ATG8 delipidation. Furthermore, we validated our observations using two artificially primed ATG8 isoforms (ATG8E ΔC and ATG8F ΔC) in two independent ATG4-loss-of function mutants (*atg4a-1/atg4b-1* and *atg4a-2/atg4b-2*, the former of which was used in the original study about the role of ATG8-delipidation in Arabidopsis ^8^).

Importantly, upon thorough evaluation, we determined that none of our results directly contradicted data presented in the study by Yoshimoto and colleagues^8^. The differences in our conclusions probably stem from serendipitous factors. For example, in our study we elected to test an artificially truncated ATG8E and ATG8F isoforms which unexpectedly turned out to be more potent in enabling autophagic flux than the natively truncated ATG8I used by Yoshimoto and colleagues. Furthermore, as our results clearly demonstrate, overexpression of GFP–ATG8I yields a lower autophagic activity in *atg4a/b* than in WT. Therefore, it is conceivable that the activity in *atg4a/b* might have fallen below the detection limit for the assay relying on the imaging of the diffuse GFP signal diluted in the vacuolar volume, implemented by Yoshimoto and colleagues. In our study we opted for imaging autophagic bodies represented by bright puncta of concentrated GFP signal, thereby increasing the chance of detecting even weak autophagic activity.

Based on our observations, we propose that plant autophagosome maturation diversified into strategies dependent on and independent from ATG4 delipidating activity (**Fig. 7**). It is an intriguing question, how a critical step regulating autophagosome maturation became unnecessary in some eukaryotes. Noteworthily, the concept of “conserved step” in this context should be viewed with some caution: while disruption of ATG8 delipidation indeed halts autophagosome biogenesis in yeast and mammalian cells, it affects different stages of the process in these organisms^6,10,12,13^, indicating inherent differences in why this step is critical for autophagy in various eukaryotes. Additionally, more recent studies suggest that the dependency of mammalian autophagy on ATG8 delipidation might be model/condition-specific^21,32^.

**Figure 7.**
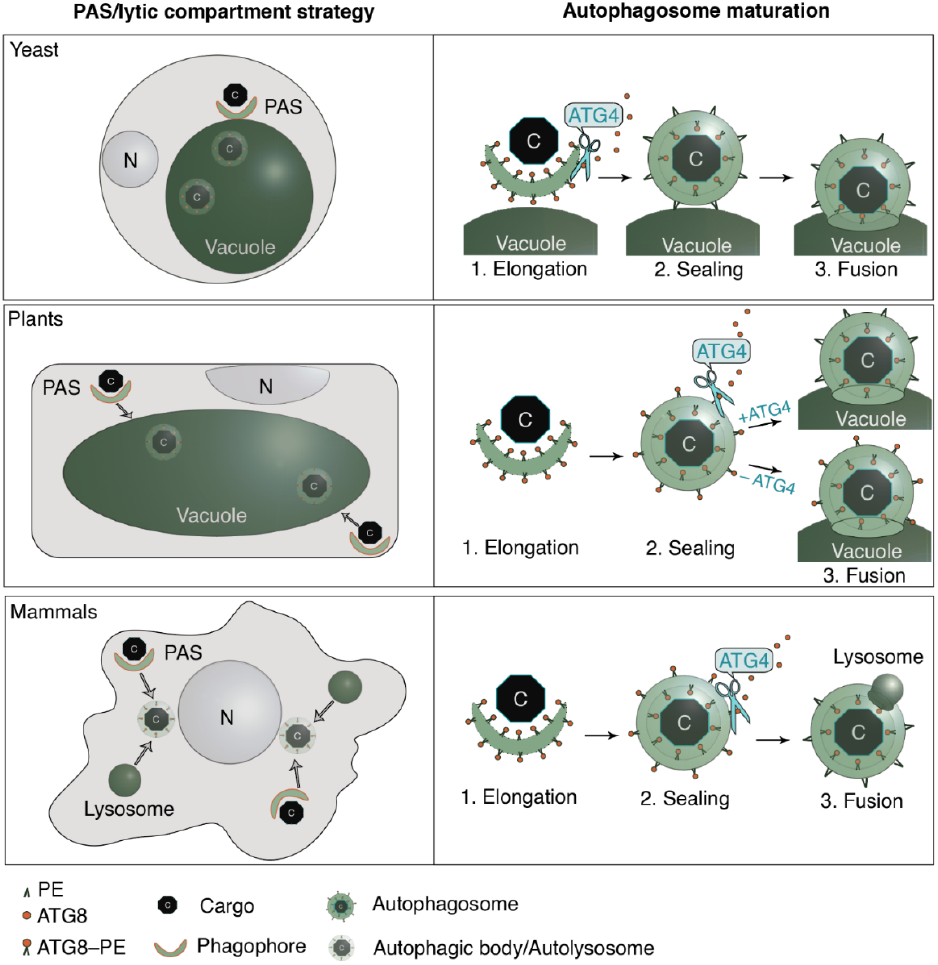
Schematic representation of ATG8 delipidation importance for autophagy in the different eukaryotic kingdoms. ATG4-dependent delipidation of ATG8 is considered to be generally conserved in plants, animals and fungi. However, this process is critical for early steps of autophagosome maturation in yeast cells (phagophore elongation), while in mammalian cells the same delipidation step is critical for the later stage of autophagy (docking of complete autophagosomes to lyso-somes). In this study we show that although ATG8 delipidation occurs in plants, it is dispensable for autophagy. The different roles ATG8-delipidation plays in autophagosome maturation might be representative of autophagy adaptation to the various strategies for the lytic compartment (single large immobile vacuoles in plant and yeast cells *vs* numerous small motile lysosomes in animal cells), for phagophore assembly sites (single site in yeast *vs* numerous sites in animal and plant cells), and autophagosome trafficking towards the lytic compartment (autophagosome formation in the proximity to the vacuole in yeast, trafficking of autophagosomes towards immobile vacuole in plants, coordinated trafficking of autophagosomes and lysosomes to their meeting spot in mammalian cells). C, cargo; N, nucleus, PE, Phosphatidylethanolamine.

Autophagy was likely invented by the Last Eukaryotic Common Ancestor (LECA)^33^ to maintain the newly acquired complex cell structures, such as organelles. During the evolution of eukaryotes, autophagy has diversified to adapt to the various life strategies, mobility traits, nutrient requirements, and specific configurations of endomembrane trafficking systems observed in fungi, animals, plants and algae. For example, eukaryotes evolved different configurations of the lytic compartment — the final destination of the autophagic pathway. The main lytic compartment of animal cells are compact and mobile lysosomes. Upon the induction of autophagy, both lysosomes and autophagosomes are trafficked towards their fusion location (**Fig. 7**). The modest size of lysosomes and their importance for other cellular pathways compels animal cells to regenerate these organelles after their fusion with autophagosomes, in a process known as autolysosome recovery (ALR)^34^. Such a process has not yet been reported for fungal and plant cells, in which autophagosomes fuse with the large lytic vacuoles that often occupy most of the cell volume^1,3^.

Notably, autophagosome biogenesis in yeast cells occurs in proximity to the vacuole, at the single phagophore assembly site^2^ (**Fig. 7**). Consequently, once complete, autophagosomes fuse immediately with vacuoles. In contrast, autophagosomes in animal cells emerge at multiple locations, and are trafficked towards nuclei to fuse there with the perinuclearly translocated lysosomes^35^. Interestingly, plants’ autophagic strategy seems to be somewhere in-between the yeast and animal solutions (**Fig. 7**). That is, autophagosomes form at multiple foci and are trafficked towards the immobile large lytic vacuole^3^. In view of these facts, it is conceivable that despite conservation of the core autophagic mechanisms, some molecular aspects of the pathway might significantly differ in vascular plants, different groups of algae, fungi and animals.

ATG8 recruits other core ATG proteins to the growing phagophore. Subsequently, delipidation of ATG8 from maturing autophagosome was suggested to release PAS proteins prior to the fusion of the autophagosome with the lytic compartment, thus preventing their unnecessary degradation^2^. Interestingly, such rescue of the core ATGs might be less critical for Arabidopsis autophagy. For instance, ATG1a was shown to be delivered to the vacuole together with autophagosomes^36^. Future studies identifying proteins decorating mature plant autophagosomes will help elucidate how they are affected by ATG8 delipidation.

In higher eukaryotes, some of the ATG gene families expanded quite significantly^3,37,38^. For example, Arabidopsis has nine ATG8 and two ATG4 orthologs. The reasons behind such expansion are not fully understood, but it is likely that functional diversification of ATG isoforms became necessary within the context of complex organisms. For instance, Zess and colleagues previously suggested a model according to which plant autophagosomes might vary in their ATG8 isoform content and consequently contain different cargo^39^. In the present study, we demonstrate drastic differences in the size of the autophagosomes formed with participation of ATG8E and ATG8I isoforms, as well as in their impact on plant stress tolerance. In the future we plan to determine the differences in the content of the ATG8F- and ATG8I-specific autophagic bodies and explain their impact on plant stress tolerance.

Current evidence indicates that Arabidopsis ATG4 orthologs are redundant: Arabidopsis knockouts of single *ATG4* genes do not have autophagy-deficient phenotypes^8,23^(**Fig. S3I** and **J**). Furthermore, we demonstrated that either of the ATG4 proteases is sufficient for complete processing of ATG8 *in planta* (**Fig. S2E** and **F**). Intriguingly, we also revealed that efficient delipidation of ATG8 requires the presence of both ATG4 proteases (**Fig. S3K** and **L**), indicating that the delipidating activity of ATG4s is significantly less efficient than their proteolytic activity, which might play a role in preserving the pool of ATG8–PE required during phagophore expansion. Similarly, highly efficient proteolytic activity and slower delipidation activity was reported for the mammalian ATG4s^12^.

Interestingly, yeast ATG4 contains the N-terminal motif, APEAR, responsible for its association with ATG8–PE during delipidation step^9^, while mammalian ATG4s were shown to associate with ATG8–PE via their C-terminal LIR motif^12^. It remains to be determined which of the predicted motives of Arabidopsis ATG4s is responsible for association with ATG8–PE. Yet, the APEAR motif is an unlikely candidate for it, as it is present only in the ATG4B, while both ATG4A and ATG4B partake in ATG8 delipidation (**Fig. S3K** and **L**).

Here we report an additional intriguing observation of naturally truncated ATG8I isoform accumulating in its ATG8I–PE adduct in the WT upon upregulation of autophagy (**Fig. 4E**). This is quite uncharacteristic for plant ATG8s, as their lipidated adducts are notoriously difficult to detect in plant protein extracts^23^. Future studies will determine whether such accumulation is caused by the decreased affinity of ATG4 towards ATG8I. Such a loss of affinity is plausible because the natively truncated ATG8 isoforms do not require interaction with ATG4 for processing before lipidation, and their interaction with ATG4 for delipidation is dispensable for autophagy.

In conclusion, we discovered that in contrast to yeast, mammals and Chlamydomonas, vascular plants do not require ATG8 delipidation for autophagosome maturation and delivery to a lytic compartment. In addition, our study provides novel insights into the isoform-specific role of ATG8 family in plant autophagy and lays foundation for future research on the mechanism of plant autophagosome maturation.

## Materials and methods

### Arabidopsis plant material and growth conditions

Unless stated otherwise, all experiments were performed in Col-0 Arabidopsis accession (N1092). Autophagy-deficient mutants were described previously: *atg4a-2/atg4b-2*^23^ (*atg4a/b* in this study), *atg5-1*^18^(*atg5* in this study), *atg7-2*^40^ (*atg7* in this study). WT and ATG4-deficient mutant (*atg4a-1/b-1*)^8^ in Was-silewskija-0 accession (N1602) were kindly provided by Prof. D. Hofius.

For cultivation on plates, Arabidopsis seeds were surface sterilized with bleach. In brief, seeds were agitated for 20 min in 10% V/V bleach (Klorin, Sweden) water solution supplemented with 0.05% Tween-20, washed in four changes of sterile MQ water and plated on 0.5xMS medium, comprising 0.5xMS (*Duchefa*, ref. M0222); 10 mM MES (*Duchefa*, ref. M1503); 1% sucrose; pH 5.8; 0.8% Plant agar (*Duchefa*, ref. P1001). Plates were sealed with plastic wrap and incubated vertically under long day conditions: 16/8-hour light /dark cycle at PAR 150 μmol photons m^−2^ s^−1^, at 22°C.

S-Jord soil (Hasselfors, Sweden) was used to grow Arabidopsis plants in pots. Plants were incubated under long day conditions: 16/8-hour light /dark cycle at PAR 150 μmol photons m^−2^ s^−1^, at 22°C.

### Chlamydomonas strains and growth conditions

UV mutagenesis strain 4 (UVM4) capable of efficiently overexpressing foreign genes^41^ was used as the WT background in this study. The *Chlamydomonas* cells were grown in tris-acetate-phosphate (TAP) medium at 23°C and a light-dark cycle (LD16:8). The cells were harvested at a density of 1-2×10^7^ cells ml^-1^ for immunoblot analysis or genomic DNA isolation.

### Cloning of constructs

Constructs for studies in Arabidopsis was generated using GreenGate^42^ and Gateway (Invitrogen) cloning systems. For expression of recombinant ATG4 proteases we created constructs coding N-terminal GST fusions of the wild-type and proteolytically dead (PD) Arabidopsis ATG4 proteins. For this, ATG4A (AT2G44140) and ATG4B (AT3G59950) CDS were lifted from total cDNA of *Arabidopsis thaliana* using primers AM 417/418 and AM 419/420, respectively (**Table S1**). Active sites of ATG4A and ATG4B were mutated using QuikChange II Site-Directed Mutagenesis Kit (Agilent, 200523) and primer pairs AM 622/AM 623 and MA 624/625, respectively (**Table S1**). Amplicons were recombined with pDONR/Zeo vector using Gateway cloning system (Invitrogen). The resulting constructs AM 753 and AM 754 (**Table S2**) further recombined with pDEST15 (Gateway™, Invitrogen) to obtain AM 810-813 destination constructs (**Table S2**).

The constructs encoding EosFP–AtATG8E variants for expression in plants were generated using Gateway cloning system (Invitrogen). To clone full-length ATG8E, ATG8E G/A and ATG8E ΔC, the genomic sequence of ATG8E (*AT2G45170*) was lifted from the total genomic DNA of *Arabidopsis thaliana* using primers AM 475/476, AM 475/478 and AM 475/477, respectively (**Table S1**). Amplicons were recombined with pDONR/Zeo vector using Gateway cloning system (Invitrogen). The resulting constructs AM 655-657 (**Table S2**) were further recombined with pUBN EosFP^43^ to obtain AM 661-663 destination clones (**Table S2**).

The construct encoding EosFP–AtATG8A– CFP for expression in plants was generated by first fusing ATG8A gene with CFP CDS using overlay PCR. For this, ATG8A gene (*AT4G21980*) was lifted from the total genomic DNA of *Arabidopsis thaliana* using primers AM 429/430, CDS of CFP was amplified from pGWB 661^44^ (**Table S1**). The obtained amplicons were fused in a PCR with primers AM 431/AM 428 (**Table S1**) and the product was recombined with pDONR/Zeo vector using Gateway cloning system (Invitrogen). The resulting constructs AM 556 (**Table S2**) was further recombined with pUBN EosFP^43^ to obtain AM 567 destination clone (**Table S2**).

The constructs encoding GFP–ATG8E ΔC, GFP–AtATG8F, GFP–AtATG8F ΔC and GFP– AtATG8I fusion proteins for expression in plants were generated using GreenGate system, following the published instructions^42^. pGGC ATGF (plasmid SH3, **Table S2**) and ATG8I (plasmid SH4, **Table S2**) modules were generated by recombining the pGGC0 module with the PCR products obtained on the total cDNA of *Arabidopsis thaliana* using primers SH PR 7/SH PR 8 and SH PR 9/SH PR 10, respectively (**Table S1**). The entry clones with ΔC versions of ATG8E and ATG8F (AM 821 and 822, **Table S2**) were generated by amplifying the sequences using primer pairs AM 611/AM 643 and SH PR7/AM 643, respectively (**Table S1**), entry clones with the full-length versions of the corresponding cDNAs were used as templates for the PCRs. pGGA 2×35S promoter (SH 117, **Table S2**) was produced by recombining the plasmid with the PCR product using primer pair SH PR 34/SH PR 35 (**Table S1**). The rest of the required modules were kindly provided by K. Schumacher’s and A. Maizel’s labs. The destination vectors AM 780, AM 783, AM 831, and AM 832 (**Table S2**) were obtained by recombining pGGZ004, SH 117, pGGB -E2GFP-GSL, the corresponding entry clone, pGGD decoy, pGGE NosT and pGGF FAST Red pLB012. AM 784 destination clone encoding free GFP driven by the 2×35S promoter (**Table S2**) was obtained by replacing the entry clones with a decoy module in the above-described recombination reaction.

SH 3 and SH 4 clones were also used to create construct expressing mScarlet-tagged AtATGF and AtATG8I, respectively (SH 17 and SH19, **Table S2**).

Cloning of constructs for experiments in *Chlamydomonas* was done using the modular cloning MoClo toolkit^45^. CrATG8 (Cre16.g689650, Phytozome accession number, https://phyto-zome-next.jgi.doe.gov/) and CrAT8 ΔC were amplified using primers ATG8_F and ATG8_R or ATG8 ΔC _R (**Table S1**) and genomic DNA as a template. Amplicons were cloned into the B5 module (L0 module) of the MoClo system.

Two L1 vectors were constructed using the following L0 modules: *PsaD* promoter (pCM0-016), *mCherry* gene (pCM0-067), the genomic sequence of *CrATG8* or *CrATG8* ΔC, and the *PsaD* terminator (pCM0-114). L2 vectors were assembled with hygromycin resistance. The L2 vectors were introduced into WT, *atg4* and *atg5* strains using the glass beads methods^46^.

### Establishment of Arabidopsis transgenic lines

Transgenic lines established for this study are listed in **Table S3**. WT, *atg4a/b* and *atg5* Arabidopsis plants were transformed using the standard floral dip method^47^ and *Agrobacterium tumefaciens* strain GV3101. Transgenic plants were selected on the corresponding herbicide or, in the case of FastRed selection marker, by detecting the red fluorescence in the seeds using an upright epifluorescence microscope (Axioplan, Zeiss). Fluorescent signal in the seedlings was verified using CLSM (LSM800, Zeiss and SP8, Leica). Expression of the protein with the expected molecular weight was confirmed by a Western blot analysis. The presence of the T-DNA inserts in the *atg4a/b* and *atg5* backgrounds was verified by genotyping (**Fig. S2A** and **B**).

To obtain single *ATG4* knockout lines expressing EosFP–ATG8E ΔC or EosFP–ATG8A– CFP, we performed crosses between *atg4a/b* plants carrying the respective constructs and WT plants carrying the same constructs.

### Establishment of Chlamydomonas knockouts using CRISPR/Cas9

A CRISPR/Cas9-based targeted insertional mutagenesis approach^48^ was used to knock out the *CrATG4 (Cre12*.*g510100*, Phytozome accession number) and *CrATG5 (Cre14*.*g630907*, Phytozome accession number) genes. For each gene, a guide RNA was designed for targeting the exon region near the 5’ UTR (**Table S1**). Electroporation was used to introduce a donor DNA comprising *Hsp70/RBCS2* promoter, *AphVIII* gene and *RBCS2* terminator, in addition to the ribonucleoprotein complex. The resulting transformants were screened with colony-PCR and confirmed by immunoblotting if specific antibodies were available.

### Transient expression in Arabidopsis protoplasts

For the experiment shown in **Fig. S7C**, protoplasts were isolated from true leaves of 1.5-month-old plants of WT, *atg4a/b* and *atg7* backgrounds. Cells were then transformed with SH 17 and SH 19 constructs (**Table S2**) to over-express mScarlet–ATG8F and mScarlet– ATG8I, respectively. Protoplast isolation, transformation and induction of autophagy was performed as previously described by Elander et al^49^. Cells were imaged using CLSM800 (Zeiss): 40x/1.2W lens, excitation light 561 nm and emission in the range of 570-650 nm. Images were processed using ImageJ software^50^ and a dedicated macro: https://github.com/Aly-onaMinina/ImageJ-macros-for-CLSM-files-procesing.

For the experiment presented in **Fig. S5** protoplasts were isolated from 4-week-old WS-0 WT and *atg4a-1/b-1* plants and transformed with the plasmids AM 779, 780, AM 831 and AM 832 (**Table S2**) to express full-length and artificially primed ATG8E and ATG8F isoforms. Cell isolation, transformation, treatment, imaging and image analysis were performed as described above.

### Genotyping of Arabidopsis lines

Genomic DNA was extracted using a quick method. In brief, approximately 5 mg of plant material was lysed in 30 μl of 0.5 M NaOH by shaking together with glass beads, pH was then equilibrated with 370 μl of 0.1 M Tris-HCl, pH 6.8. Three μl of the resulting DNA solution was used for each PCR reaction.

*atg4a-2* mutation was verified using primer pairs annealing on the WT *ATG4A* allele (AM 409/AM 587, **Table S1**) and on the *atg4a-2* T-DNA insertion (AM 410/AM 584, **Table S1**). *atg4b-2* mutation was verified using primer pairs annealing on the wild-type *ATG4B* allele (AM 423/AM 420, **Table S1**) and on the *atg4b-2* T-DNA insertion (AM 420/AM 584, **Table S1**). Detection of the *atg5-1* T-DNA insertion was done using primers AM 1/AM 579 (**Table S1**), while WT ATG5 gene was detected with primers AM 1/AM 2 (**Table S1**).

### Isolation of the Chlamydomonas Genomic DNA

Genomic DNA was isolated as previously described^51^. The cells were grown under standard conditions and harvested at a density of 1-2×10^7^ cells ml^-1^.

### RT-PCR

Total RNA was extracted from 1.5-week-old Arabidopsis seedlings pooling 3 seedlings/extraction for each transgenic lines and using Spectrum™ Plant Total RNA Kit (Sigma-Aldrich, STRN250), RNAs were on-column treated with the DNase (Sigma-Aldrich, DNASE70-1SET) following the kit instructions. One μg of total RNA was used for each RT reaction implementing Maxima First Strand cDNA Synthesis Kit (ThermoFisher, K1641) and oligodT primers provided with the kit, and following the manufacturer’s protocol. Full length cDNAs of the ATG4A, ATG4B and WT were detected using primer pairs AM 417/AM 418, AM 419/AM 420, and AM 8/AM 9, respectively (**Table S1**). The amplicons were separated in a 1% TAE agarose gel containing GelRed dye and imaged with ChemiDoc (Bio-Rad).

### qPCR

Primers for ATG8 isoforms were designed and checked for specificity against Arabidopsis transcriptome using Primer-Blast (**Table S1**). Primers for reference genes were taken from the previous study^52^. Total RNA extraction and RT were performed as described above, at least 15 seedlings were pooled for each line for the RNA extraction. RNA RIN was confirmed to be above 8. 1000 ng of total RNA were used for each cDNA synthesis reaction, RT reactions were diluted 1:25 and 5 μl were used for each qPCR reaction (final volume of 15 μl), each reaction was run in triplicates. qPCR was performed using the DyNAmo Flash SYBR Green qPCR Kit (ThermoFisher, F-415L), the recommended in the manual two-step protocol supplemented with the melt curve step. Primers were used at the 0.5 μM final concentration and annealed at 60°C. The assay was carried out in the CFX Connect (Bio-Rad) and analyzed using CFX Maestro software (Bio-Rad). The lack of the off-target amplification was verified by the melt curve analysis and stability of the reference genes was confirmed by ideal M-values identified by the software. Expression of each GOI was normalized to all three reference genes using the ΔΔCt method and presented as the fold change relative the WT sample devoid of any transgenes.

### In planta ATG8-CFP cleavage assay

True leaves of one-month-old Arabidopsis plants expressing EosFP–ATG8A–CFP in *atg4a, atg4b, atg4a/b* and WT backgrounds were harvested and immediately flash-frozen in liquid nitrogen. Plant material was powdered while frozen and mixed 1:2 (m:V) with pre-heated 2xLaemmli buffer. Samples were boiled for 10 min at 95 °C and centrifugated at 17,000 *g* for 10 min. Seven and a half μl of each supernatant were loaded on a 4-20% TGX Stain-Free precast gel (Bio-Rad, 4568096). Separated proteins were transferred onto PVDF membrane (Bio-Rad, 1620174) and blotted with anti-GFP (Roche, 11814460001) and anti-EosFP (Encor-bio, RPCA-EosFP), both at 1:2,000 dilution. Anti-mouse (Amersham, NA931-1ML) and anti-rabbit (Invitrogen, a11008) HRP conjugates were used at 1:5,000 and 1:20,000 dilutions, respectively. Signal was developed using SuperSignal™ West Femto Maximum Sensitivity Substrate (ThermoFisher, 34094) and Chemi-Doc Imaging System (Bio-Rad). Densitometric analysis was performed as described in the dedicated chapter below.

### In vitro ATG8-CFP cleavage assay

AM 810–813 plasmids (**Table S2**) encoding active and proteolytically dead (PD) GST-tagged AtATG4A and AtATG4B were used to transform Rosetta (DE3) cells (Novagen, 70954-3). Expression was induced overnight at 16°C using 0.25 mM IPTG (Hellobio, HB3941). Cell cultures were harvested by centrifugation at 5,000 *g* for 20 min, and the cell pellets were stored at –80°C until purification. Subsequent steps were conducted at 4°C. Cells were resuspended using a ratio of 1 g to 5 ml of binding buffer containing 20 mM PBS buffer, 100 mM NaCl, at pH 7.3. Cells were lysed using a cell disruptor at 20,000 psi. The resulting sample was then treated with DNase I for 30 min. Insoluble cell debris was eliminated through centrifugation at 38,000 *g* for 20 min, followed by filtration of the supernatant through a 0.45 μm filter. The filtered supernatant was applied to an 8.5 mm diameter column packed with 1 ml of Glutathione Sepharose 4B matrix (Cytiva, 17075601) and equilibrated with the binding buffer. After sample application, the column was washed with approximately 10 column volumes of the binding buffer. The protein was eluted using an elution buffer containing 50 mM Tris-HCl, 100 mM NaCl, 10 mM reduced glutathione, at pH 8.0. Each fraction was collected after a 5-min incubation with the elution buffer. The protein purity was assessed on 4– 20% gradient SDS-PAGE gels. The purest fractions were combined and concentrated to 1 ml using Vivaspin 30K centrifugal concentrators (Sartorius, VS0621). The protein sample was then further purified by size exclusion chromatography on a HiLoad 16/600 Superdex 200 pg column (Cytiva, 28989335) connected to an ÄKTA FPLC system (Cytiva). The column was equilibrated with a buffer containing 50 mM Tris-HCl, 100 mM NaCl, 1 mM DTT at pH 8.0. Protein fractions were analyzed with SDS-PAGE, pooled and concentrated to desired concentration. The protein concentration was calculated based on the measured A280 nm using a NanoDrop 1000 Spectrophotometer and the theoretical molar extinction coefficient determined using the ProtParam tool on the ExPASy server.^53^ Before storing at –80°C, protein sample was mixed with glycerol in a 1:1 ratio. True leaves of one-month-old Arabidopsis plants expressing EosFP–ATG8A-CFP in *atg4a/b* and WT backgrounds were harvested and immediately flash-frozen in liquid nitrogen. Plant material was powdered while frozen and mixed 1:2 (m: V) with reaction buffer containing 50 mM Tris pH 7.5, 100 mM NaCl, 5 mM DTT, 5 mM EDTA, 5 mM PMSF, 0.05% Tween-20 and 50 ng of purified active or PD ATG4. Reactions were kept at room temperature for the indicated period of time, prior to be mixed 1:1 with preheated 2xLaemmli buffer. Detections of EosFP and CFP in the reaction were preformed using Western blot assay as described for the *in planta* ATG8-CFP cleavage assay. Densitometric analysis was performed as described in the dedicated chapter below.

### In vitro ATG8 delipidation assay

True leaves of one-month-old Arabidopsis plants expressing EosFP–ATG8E ΔC in *atg4a/b* and WT backgrounds were harvested and immediately flash-frozen in liquid nitrogen. Plant material was powdered while frozen and mixed 1:2 (m: V) with reaction buffer containing 50 mM Tris pH 7.5, 100 mM NaCl, 5 mM DTT, 5 mM EDTA, 5 mM PMSF, 0.05% Tween-20 and 50 ng of purified active or PD GST-ATG4. Reactions were kept at room temperature for 30 min, prior to being mixed 1:1 with preheated 2xLaemmli buffer. To estimate the original amount of ATG8 and ATG8-PE present in the samples, a quick protein extraction was performed by mixing an aliquot of frozen plant material directly with preheated 2xLaemmli buffer at 1:1 m:V ratio. Proteins from the quick extracts and delipidation reactions were separated on the manually casted 10% PAAG supplemented with 6 M urea^23,54^. To ensure sufficient separation of the tagged ATG8 adducts, the electrophoresis was performed until the 25 kDa band of the prestained PageRuler (ThermoFisher Scientific, 26619) reached the bottom edge of the gel. Separated proteins were transferred onto PVDF membrane (Bio-Rad, 1620174). Detection of EosFP was performed as described for the *in planta* ATG8-CFP cleavage assay. Densitometric analysis was performed as described in the dedicated chapter below.

### Autophagy induction

Induction of autophagy in Arabidopsis seedlings was performed in liquid 0.5xMS medium complemented with 5 μM AZD8055 (Santa-Cruz Biotech, 364424) and 0.5 μM concanamycin A (Santa-Cruz Biotech, 202111A). Treatment for 2 h was sufficient for induction of autophagy in seedlings roots, while treatment for 24 h was used for cotyledons.

For autophagy induction in older plants, the abaxial side of true leaves were infiltrated with MQ water containing 5 μM AZD and 0.5 μM ConA. The adaxial side of the infiltrated leaves was imaged after 24 h of treatment.

Treatments with 10 mM N-ethylmaleimide (NEM, Sigma-Aldrich 04259) and 1 mM iodoacetamide (IAM, Cytiva RPN6302) were performed simultaneously with AZD and ConA due to high phytotoxicity of the NEM and IAM. These treatments were performed using the approach described above for AZD/ConA. E64d showed lower toxicity in Arabidopsis, therefore plant material was pretreated with 5 μM E64d (Sigma-Aldrich E8640) prior to autophagy induction with AZD/ConA.

Nitrogen depleted conditions were implemented by growing the seedlings starting from seed germination on 0.5xMS devoid of nitrogen source: 0.5xMS basal salt mix (Sigma-Aldrich M0529): 1.5 mM CaCl_2_, 0.75 mM MgSO_4_, 0.25 mM KH_2_PO_4_, 2.5 mM KCl, 1% sucrose and 0.8% Plant agar.

Carbon depleted conditions were achieved by using 0.5xMS (described above) without sucrose and keeping plants in the dark.

### Confocal microscopy

Confocal microscopy was preformed using CSLM SP8 (Leica) and CLSM 800 (Zeiss), in either case using 40x water immersion lens, NA=10–1.2. GFP, CFP and EosFP were imaged using excitation light of 488 nm and emission range 515–560 nm. Red fluorescence was detected using 561 nm excitation light and 570– 650 nm emission.

To prevent root desiccation during imaging, seedlings were mounted inside RoPod chambers^55^, in a droplet of liquid corresponding to the treatment. Detached cotyledons were imaged while mounted between two coverslips. Ten by ten mm sections were cut out of true leaves immediately prior to imaging and mounted in a droplet of liquid between standard microscopy slide and a coverslip additionally secured with tape. True leaves and cotyledons were imaged only from the adaxial side.

Confocal images were processed using LAS X (for Leica micrographs) software or ImageJ^50^ implementing the dedicated macro for channel separation and image cropping: https://github.com/AlyonaMinina/ImageJ-macros-for-CLSM-files-procesing.

### Quantification of autophagic bodies density and area

Quantification of autophagic bodies was performed in a semi-automated manner using ImageJ software and custom-made macro. The puncta quantification shown in **Figs. 2, S3** and **S4** was done using the macro for puncta quantification in single channel v4. It can be found in the dedicated GitHub repository: https://github.com/AlyonaMinina/Puncta-quantification-with-ImageJ.

The quantifications shown in **Figs. 4** and **S9** were obtained using macro for autophagic body size and density measurement: https://github.com/AlyonaMinina/Autophagic-bodies-size-measurement..

Statistical analysis of the quantitative data obtained with ImageJ macros was performed using custom R scripts. Unless stated otherwise, results were subjected to Tukey’s HSD test. Plots were built using the ggplot2 R package^56^. Each experiment was performed at least twice, using at least three biological replicates (individual plants), three technical replicates for imaging (images of representative areas on the same plant) and five to ten technical replicates for measurements (regions of interest selected on each image using the corresponding macro script).

### TEM

Seven-day-old seedlings were treated with 5 μM AZD and 0.5 μM ConA, in 0.5x MS liquid medium, subjected to confocal microscopy and then fixed in the solution containing 2.5% glutaraldehyde and 1% paraformaldehyde in 0.1 M phosphate buffer. Further sample processing was performed at BioVis, Uppsala: the samples were post-fixed with 1% Osmium tetroxide, dehydrated in ethanol embedded in resin. Ultrathin sections were collected on copper slot grids and contrasted with uranyl acetate and lead citrate. Sections were imaged using FEI Technai G2 operated at 80 kV. Images were processed using ImageJ 1.53c.

### Western blotting

Whole Arabidopsis seedlings or a section of true leaf were sampled into liquid nitrogen and powdered while frozen. A pool of 4–8 seedlings per sample or sections from three different leaves were used as one sample to ensure representativity of the data. To limit potential protein proteolysis during sample preparation, the powdered material was directly mixed with pre-heated Laemmli buffer in 1:2 (m: V) ratio. Samples were boiled at 95°C for 10 min, mixing every 2 min. Debris was removed by centrifugation at 17,000 *g* for 10 min. Supernatants were transferred into new Eppendorf tubes and stored at –20°C. Typically, 7.5-15 μl of each sample were used for separation on the gel.

To separate tagged ATG8 and tagged ATG8–PE adducts, 10% PAAG was prepared manually supplementing it with 6 M urea^23,54^. To ensure sufficient separation of the tagged ATG8 adducts, the electrophoresis was performed until the 25 kDa band of the prestained PageRuler (ThermoFisher Scientific, 26619) reached the bottom edge of the gel. For other Western blot assays, proteins were separated on 4–20% TGX Stain-Free precast gels (Bio-Rad, 4568096). Detections of the EosFP, GFP and CFP were performed using Western blot assay as described for the *in planta* ATG8–CFP cleavage assay.

Total proteins from *Chlamydomonas* cells were extracted as described in Zou et al., (2024)^51^. The cells were grown under standard conditions and harvested at a density of 1-2×10^7^ cells ml^-1^. Anti-CrATG4 (Agrisera, AS15 2831) was used at dilution 1:5,000, while anti-rabbit HRP-conjugate (Agrisera, AS07 260) at a dilution of 1:10,000.

### Densitometric analysis

Densitometric analysis of the western blots was performed using custom-designed ImageJ macro. GFP-cleavage assay macro was used for EosFP/GFP-cleavage assay shown in **Fig. 2E, Fig. 4D, Fig S4D** and **Fig. S8D**. The script and instructions for users are available on GitHub: https://github.com/AlyonaMinina/NACA-N-terminal-ATG8-cleavage-assay.

Densitometric analysis for the C-terminal ATG8 cleavage assay shown in the **Fig. S2E** and **G** were carried out using the dedicated ImageJ macro (https://github.com/AlyonaM-inina/CACA-C-terminal-ATG8-cleavage-assay).

For the **Figs. 2D, S3G, S3K and S3L** ATG8-lipidation was quantified using the ALDA macro version for western blots with two bands; while quantification for **Fig. 4E** was made using the macro version for western blots with three bands. Both ImageJ macros are available on GitHub: https://github.com/AlyonaM-inina/ALDA-ATG8-lipidation-densitometric-assay.

Quantitative ImageJ macro outputs were further processed in R Studio using custom R scripts for sample relabeling and Tukey’s HSD test, plots were built using the ggplot2 R package^56^.

### Plant phenotyping

Root growth assays were performed using SPIRO^24^ for automated imaging of Petri plates and SPIRO assays for image analysis. To ensure reproducibility of the conditions and therefore comparability of the results, SPIRO robots were placed inside the same plant growth cabinets that were used for growing seedling on plates in other experiments of this study. For experiments illustrated in **Fig. 3, Movies S2** and **S3**, seeds were plated directly on the corresponding medium used for the stress induction and plates were imaged without interruption during day and night for at least 7 days, utilizing green LED of SPIRO for imaging in the dark. For carbon starvation assays, seedlings were allowed to grow under long day conditions for 4– 5 days, followed by 4 days of dark treatment and another 4 days of recovery.

For the experiments shown in **Fig. 4I** and **4J** seeds of the T2 generation, showing red fluorescent signal of the FastRed marker, were first germinated on standard 0.5xMS medium. Roots of 4-day-old seedlings were checked with a stereoscope to ensure strong and even GFP signal. Seedlings with confirmed transgene expression were then transferred onto –N medium and imaged using SPIRO for 7 days.

SPIRO data analysis was performed as described by Ohlsson et al ^24^.

Photos of plants in soil were obtained using the camera of an iPhone XI and further processed in Photoshop 25.0.0.

### Senescence assay in Chlamydomonas

The *Chlamydomonas* cells were pre-cultured until they reached the early stationary phase. The cells were then subjected to ten times dilution using fresh TAP medium. Upon attaining the late log phase or stationary phase, 1x10^5^ cells were inoculated on TAP agar plates for senescence assay. Cell colonies were imaged after 40 days.

### Protein sequence alignment and phylogenetic tree building

Protein sequence alignments and construction of the phylogenetic tree was done using the Geneious software v10.2.6.

## Data availability

Raw imaging data and SPIRO time-lapse imaging data will be provided by the corresponding author upon request.

## Authors contributions

S.H. contributed to analysis of experiments and editing of figures and the manuscript; Y.Z. performed all experiments involving *Chlamydomonas*; J.A.O. contributed to analysis of experiments and editing of figures and the manuscript; I.S. contributed to analysis of experiments and the manuscript editing; J.X.L. contributed analysis of experiments and the manuscript editing; F.B. contributed analysis of experiments and the manuscript editing; M.K., K.S., S.U. and Y.D. provided unpublished material and contributed to the manuscript editing;, P.N.M, S.S and P.V.B. contributed editing of figures and the manuscript; E.A.M. contributed the study concept; planning, performance and analysis of experiments; figure preparation and writing of the manuscript.

### Acknowledgments

We would like to thank Monika Hodik and Karin Staxäng at EM facility, BioVis, Uppsala for help with the TEM experiments.

This study was supported by grants from EU Horizon 2020 MSCA IF (799433) and Carl Tryggers Foundation (CTS 14 326 and 20:287) to EA Minina; The Swedish Research Council Formas (2017-00541 to PV Bozhkov and 2021-01812 to EA Minina); The Swedish Research Council Vetenskapsrådet (2019-04250 to PV Bozhkov, EA Minina, and PN Moschou); The Knut and Alice Wallenberg Foundation (2018.0026 to PV Bozhkov and PN Moschou; 2021.0071 to S Stael), European Research Council 2021 Consolidator (101044878 to S Stael), SFB1101 (TPA02, DFG to K Schumacher); Austrian Science Fund (FWF-P34944 to Y. Dagdas); European Research Council Grant (101043370 to Y. Dagdas and 948996 to S. Uestuen); Emmy Noether Fellowship GZ: UE 188/2 (DFG to S. Uestuen) and Crops for the Future Research Program at the Swedish University of Agricultural Sciences (to PV Bozhkov and to EA Minina).

## Competing interests

The authors declare no competing interests.

## List of supplementary material

### Supplementary figures and movies

**Figure S1.**
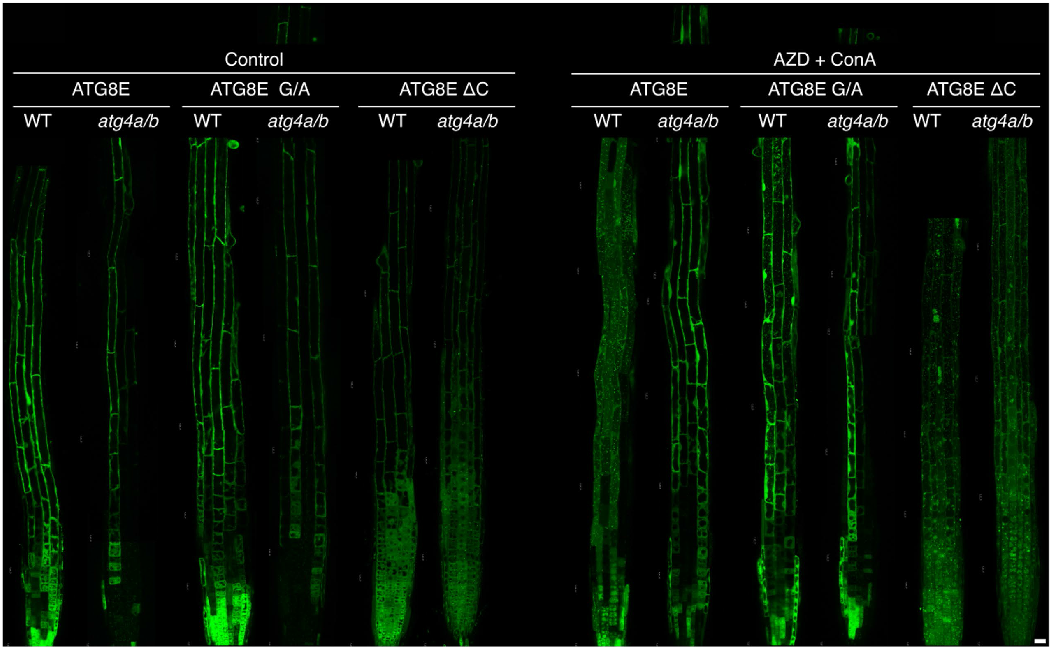
Expression of ATG8E ΔC restores accumulation of puncta in the vacuoles of ATG4-deficeint Arabidopsis roots. Confocal microscopy images showing complete tile-scans of roots partially shown in **Fig. 1 D**. Control, roots incubated and imaged in the standard 0.5xMS growth medium; AZD + ConA, roots incubated and imaged in the 0.5xMS medium supplemented with 5 μM AZD8055 and 0.5 μM concanamycin A. Scale bar, 20 μm.

**Figure S2.**
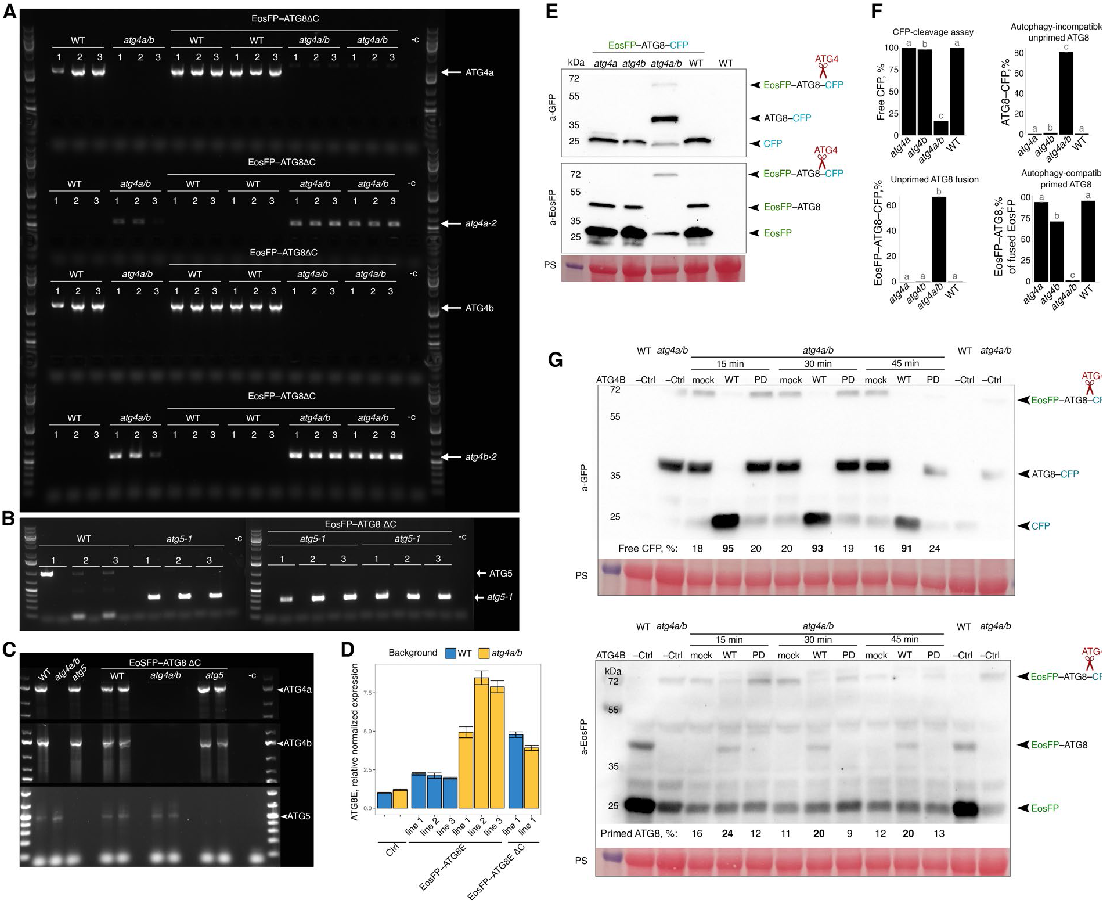
Verification of the *atg4a/b* knockout and EosFP-ATG8E ΔC -expressing lines used in the study. **A**. Genotyping of Arabidopsis lines to detect WT *ATG4A* and *ATG4B* alleles as well as *atg4a-2* and *atg4b-2* T-DNA insertions. The results confirm absence of complete *ATG4* genes in all lines with *atg4a/b* background. **B**. Genotyping of Arabidopsis lines to detect WT *ATG5* allele as well as *atg5-1* T-DNA insertion. **C**. RT-PCR confirms absence of full length *ATG4* mRNAs in the *atg4a/b* lines. Similarly, *atg5-1* lines do not contain full length *ATG5* transcript. **D**. qPCR analysis of ATG8E expression in WT and *atg4a/b* lines with and without additional transgenes. Expression was normalized to the ATG8E level detected in WT and three reference genes. Error bars show SEM for the technical triplicates. **E**. Lack of ATG4 proteolytic activity in the *atg4a/b* double knockout was confirmed by Western blot detection of the EosFP– ATG8–CFP fusion protein in the total protein extracts of plants stably expressing the fusion in the WT, *atg4a, atg4b*, and *atg4a/b* backgrounds. The product of cleavage by ATG4, EosFP–ATG8, was not detectable in the *atg4a/b* knockout line. Notably, expression of either of the ATG4 proteases is sufficient to process the ectopically overexpressed EosFP–ATG8– CFP fusion protein. PS, Ponceau S staining used as loading control. **F**. Densitometry results obtained on the Western blot shown in (**E**). Absence of both ATG4 proteases led to a drastic decrease in accumulation of free CFP (top left chart) and a corresponding accumulation of the uncut EosFP-ATG8-CFP fusion (bottom left chart). Importantly, only in the absence of both ATG4 proteases autophagy-incompatible ATG8-CFP was accumulating (top right chart), while accumulation of autophagy-competent primed EosFP-ATG8 form was dramatically decreased in the plants lacking both ATG4s. In summary, these results confirm the lack of autophagy-relevant ATG4 activity in the *atg4a/b* knockout plants. Error bars show SEM of technical replicates. Distinct letters indicate groups that are significantly different (α = 0.05), Tukey’s HSD test, *n* = 12. **G**. Processing of the EosFP–ATG8–CFP fusion by ATG4 protease was corroborated in the *in vitro* assay. Total protein extracts of plants stably expressing EosFP–ATG8–CFP fusion in *atg4a/b* background were incubated together with a buffer (mock), recombinant active ATG4B (WT) or its proteolytically dead mutant (PD) for 15, 30, or 45 min. The reactions were separated on the polyacrylamide gel and subjected to Western blotting detecting EosFP tag. Additionally, total protein extracts of plants expressing the same fusion protein in the WT and ATG4-deficient background were mixed directly with loading buffer and used as additional controls for fusion protein stability *in vivo* (WT –Ctrl and *atg4a/b* –Ctrl, respectively). Cleavage of the bond between ATG8 and CFP was observed only in the presence of active ATG4. Numbers under Western blot lanes indicate densitometry results expressed as integrated density of the indicated bands expressed as % of the total signal intensity detected in the corresponding sample. PS, Ponceau S staining used as loading control.

**Figure S3.**
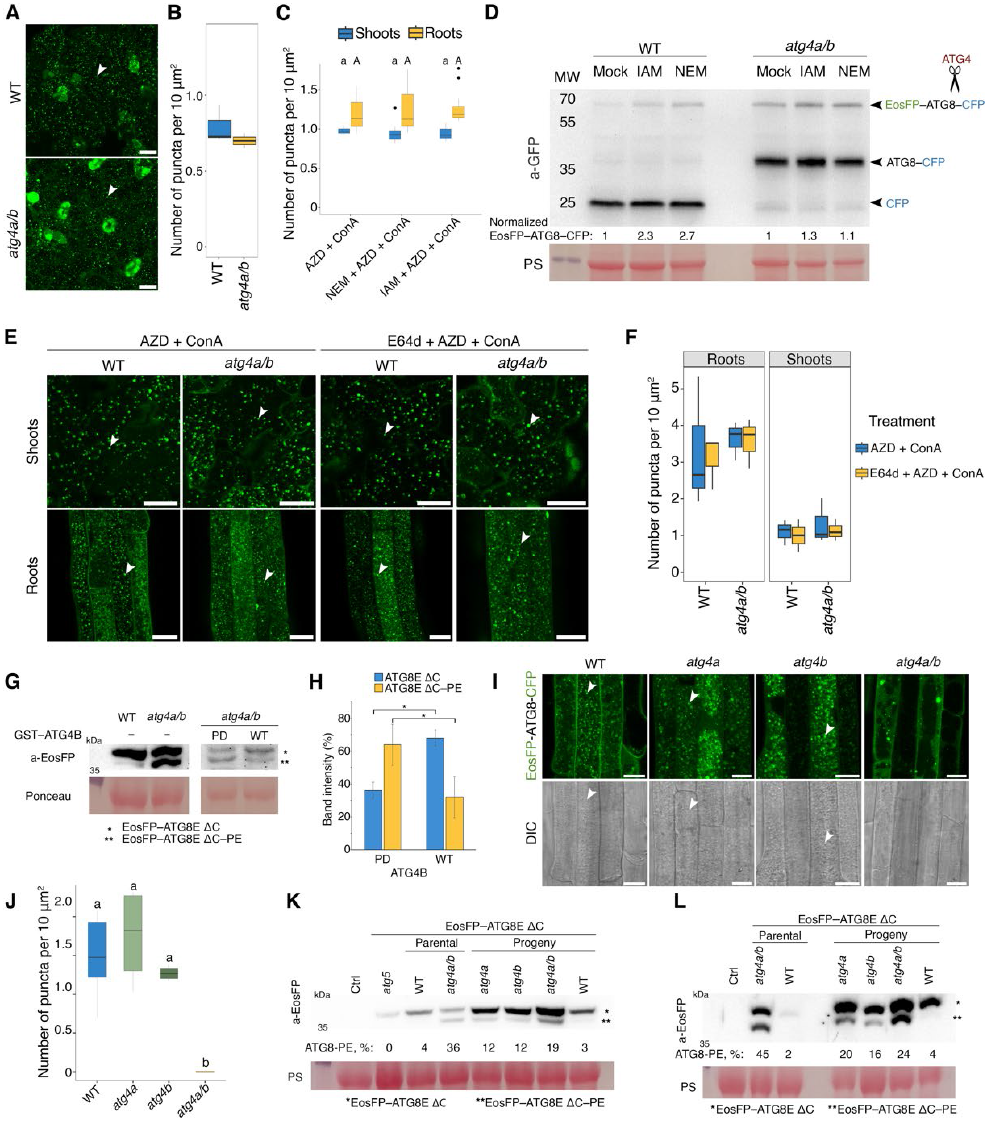
The proteolytic activity of either of the ATG4 isoforms is sufficient for autophagy, however both proteases are required for efficient delipidation of ATG8. **A**. Confocal microscopy images of leaf epidermal cells of 6-week-old Arabidopsis plants expressing EosFP–ATG8E ΔC in WT or ATG4-deficient backgrounds. Leaves were infiltrated with AZD/ConA solution 24 h prior to imaging. Scale bars, 20 μm. **B**. Quantification of puncta per vacuolar area for the samples illustrated in (**A)**. The chart shows representative results of one out of two experiments, *n* = 6 biological replicates (75 technical replicates). The samples differ according to Welch’s *t*-test, *, *p* < 0.05. Boxes indicate interquartile range (IQR); median is indicated by a horizontal line; whiskers represent range within 1.5 IQR. **C**. Quantification of puncta per vacuolar area accumulating upon NEM and IAM treatment, for the samples illustrated in **Fig. 2C**. The chart shows representative results of one out of two experiments, *n* = 61. Tukey’s HSD test was performed for each plant organ separately, *n* = 9 biological replicates (270 technical replicates), *p* < 0.05, test results for shoots are indicated with lower case letters and for roots with capital letters. Boxes indicate interquartile range (IQR); median is indicated by a horizontal line; whiskers represent range within 1.5 IQR; outliers are represented by filled circles. **D**. Western blot detection of *in vivo* EosFP-ATG8-CFP protein cleavage in seedlings treated with NEM and IAM. Seven-day-old Arabidopsis seedlings expressing the fusion protein in WT or ATG4-deficient background were treated with AZD/ConA supplemented with IAM or NEM for 24h prior to total protein extract isolation. Western blot demonstrates accumulation of the uncleaved full-length fusion EosFP-ATG8-CFP upon IAM or NEM treatment in the WT, but not in the atg4a/b plants, thereby confirming the inhibitory effect of these drug treatments on the ATG4 activity. Densitometry results are shown as numbers under corresponding lanes, representing integrated density of the uncleaved protein band expressed as % of total signal intensity for the corresponding sample. PS, Ponceau S staining used as a loading control. **E**. Confocal microscopy images of root and shoot epidermal cells of 7-day-old Arabidopsis seedlings expressing EosFP– ATG8E ΔC in WT or *atg4a/b* backgrounds. Seedlings were treated for 8 h with the cysteine protease inhibitor E64d prior to induction of autophagy with AZD/ConA. Scale bars, 20 μm. **F**. Quantification of puncta per vacuolar area for the data illustrated in (**E)**. Tukey’s HSD test was performed for each plant organ individually and did not reveal any statistically significant difference, α = 0.05, *n* = 12 biological replicates (360 technical replicates). Boxes indicate interquartile range (IQR); median is indicated by a horizontal line; whiskers represent range within 1.5 IQR. The chart shows representative results for one out of two experiments. **G**. *In vitro* ATG8 delipidation assay. Left panels: higher accumulation of the ATG8–PE adduct in *atg4a/b* compared to WT, expressing EosFP–ATG8E ΔC, was confirmed by immunodetection of EosFP in total protein extracts obtained by mixing plant material with hot Laemmli buffer. Right panels: total protein extract of *atg4a/b* plant expressing EosFP–ATG8E ΔC was incubated with recombinant active GST–ATG4B (WT) or its proteolytically dead mutant (PD) prior to Western blot detection of the EosFP tag. Ponceau S staining was used as a loading control. **H**. Quantification of the EosFP–ATG8E ΔC and EosFP– ATG8E ΔC–PE bands intensities illustrated in **G** shows significant decrease in the ATG8–PE content in the presence of active ATG4B. Band intensity is expressed as a percentage of the cumulative intensity for both bands in each sample. Error bars indicate SD. Chart shows results for two independent experiments. Student’s t-test, α = 0.05; *, p <0.001, *n* = 24. **I**. Confocal microscopy images of root epidermal cells expressing EosFP–ATG8–CFP in WT, ATG4A-, ATG4B- and ATG4A/B-deficient backgrounds. Seven-day-old seedlings were treated with AZD/ConA for 24 h prior to imaging. Accumulation of autophagic bodies is inhibited only in the absence of both ATG4 isoforms. DIC, differential interference contrast microscopy which allows visualizing autophagic bodies without labelling. **J**. Quantification of puncta per vacuolar area for the samples illustrated in (**I**) indicates very similar autophagic activities in WT and single ATG4A- and ATG4B-knockouts, while no autophagic activity was detectable in the double *atga4a/b* knock-out. These findings support the conclusion that the proteolytic activity of a single ATG4 protease is sufficient for a normal autophagic response in Arabidopsis. Distinct letters represent statistically different groups, Tukey’s HSD test, *p* < 0.05, *n* = 63 biological replicates (300 technical replicates). Boxes indicate interquartile range (IQR); median is indicated by a horizontal line; whiskers represent range within 1.5 IQR. **K**. Western blot detection of EosFP tag in the total protein extract of 14-day-old Arabidopsis seedlings expressing EosFP– ATG8E ΔC in the WT, ATG5-, ATG4A-, ATG4B-, and ATG4A/B-deficient backgrounds. The sampling was done without any prior treatment. –Ctrl, negative control (extract of Arabidopsis seedlings not expressing EosFP tagged proteins). Single *ATG4*-knockout plants were obtained by crossing *atg4a/b* plants with WT. Parental, plants grown from seed used for crossing; progeny, plants grown from F2 generation seeds obtained from the crosses. Seedlings were genotyped prior to protein extraction to confirm the genetic background. Densitometry results are shown as numbers under corresponding lanes, representing integrated density of the lipidated ATG8 band expressed as % of total signal intensity for the corresponding sample. PS, Ponceau S staining used as a loading control. **L**. Detection of the EosFP–ATG8–PE form in the total protein extracts from 6-week-old plants corroborates observations made on seedlings and illustrated in **A**: both ATG4 proteases are required for efficient delipidation of ATG8E ΔC. Sampling was done without any prior treatment. Densitometry results are shown as numbers under corresponding lanes, representing integrated density of the lipidated ATG8 band expressed as % of total signal intensity for the corresponding sample. PS, Ponceau S staining used as a loading control. White arrowheads point at autophagic bodies in the vacuoles.

**Figure S4.**
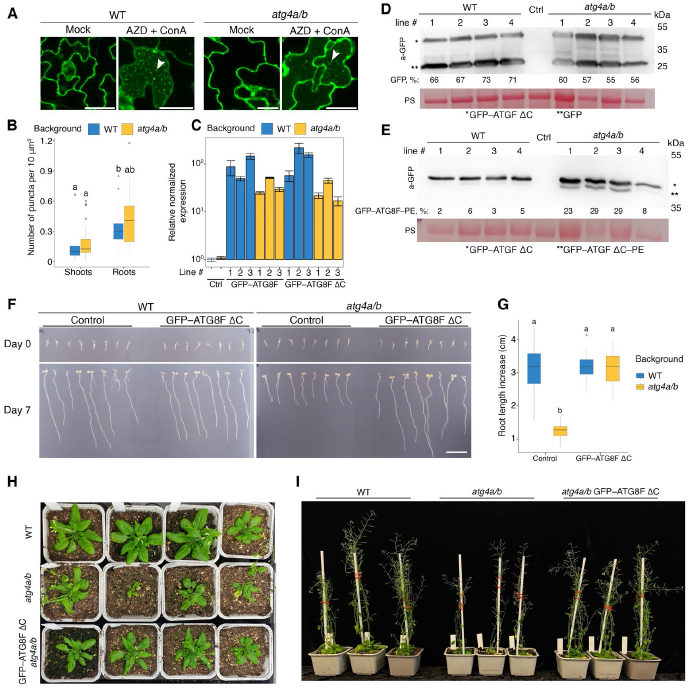
Expression of artificially primed ATG8F restores autophagic activity in ATG4-deficient background. **A**. Confocal microscopy images of leaf epidermal cells of 4-week-old Arabidopsis plants expressing EosFP–ATG8F ΔC in WT or ATG4-deficient backgrounds. Leaves were infiltrated with AZD/ConA solution or water (mock) 24 h prior to imaging. White arrowheads point at autophagic bodies accumulating in the vacuoles of treated leaves. Scale bars, 20 μm. **B**. Quantification of puncta per vacuolar area for the samples illustrated in **A** and **Fig. 2I**. The chart shows representative results of one out of two experiments, *n* = 100. Distinct letters represent groups that are significantly different from each other, Tukey’s HSD test, α = 0.05. Boxes indicate interquartile range (IQR); median is indicated by a horizontal line; whiskers represent range within 1.5 IQR; outliers are represented by empty circles. **C**. qPCR analysis of ATG8F expression in WT and *atg4a/b* lines with and without additional transgenes. Expression was normalized to the ATG8F level detected in WT and to three reference genes. Error bars show SEM for the technical triplicates. Three individual lines were checked for each genotype. **D**. GFP-cleavage assay confirms restored autophagic activity in *atg4a/b* plants expressing the GFP-tagged primed version of ATG8F (GFP–ATG8F ΔC). True leaves of 6-week-old Arabidopsis plants were infiltrated with AZD/ConA 24h prior to total protein extract isolation. Four individual lines were checked for WT and *atg4a/b* background. Densitometry results are shown as numbers under corresponding lanes and represent integrated density of the lipidated ATG8 band expressed as % of total signal intensity for the corresponding sample. PS, Ponceau S was used as a loading control. **E**. Western blot detection of the lipidated ATG8 form. Protein extract depicted in (**D**) were separated on a polyacrylamide gel containing 6M urea to reveal the accumulation of the lipidated form of ATG8 (GFP–ATG8F ΔC–PE) in the absence of ATG4 activity. Densitometry results are shown as numbers under corresponding lanes. Ponceau S was used as a loading control. **F**. Root phenotype of 12-day-old seedlings incubated for 7 days on nitrogen-depleted medium. Expression of the artificially primed ATG8F ΔC isoform is sufficient to alleviate the autophagy-deficient phenotype of *atg4a/b*. Seedlings were grown for 4 days on standard 0.5xMS medium, checked for GFP signal, then transferred onto –N medium plates to be imaged for 7 days. Scale bar, 1cm. **G**. Quantification of the increase in root length of the seedlings after 7 days of growth on the –N medium (illustrated in **F**). Boxes indicate interquartile range (IQR); median is indicated by a horizontal line; whiskers represent range within 1.5 IQR; outliers are represented by empty circles. The chart shows representative data from one experiment. Distinct letters represent groups that are significantly different from each other, Tukey’s HSD test, α = 0.05, *n* =47. Each seedling of WT or *atg4a/b* expressing GFP–ATG8F ΔC represents an independent transgenic line. **H**. Rosette phenotype of 5-week-old plants grown under long day conditions show a typical attribute for autophagy-deficiency: an early onset of senescence in the *atg4a/b background*, and lack thereof in both the WT and *atg4a/b* plants expressing ATG8F ΔC protein. Each *atg4a/b* GFP–ATG8F ΔC plant is an individual transgenic line. The experiment was performed twice using four biological replicates for each repetition **I**. Phenotypes of two-month-old plants grown in soil under long day conditions. Photos show plants with representative phenotypes, each *atg4a/b* GFP–ATG8F ΔC plant is an individual transgenic line, the experiment was performed twice using four biological replicates for each repetition.

**Figure S5.**
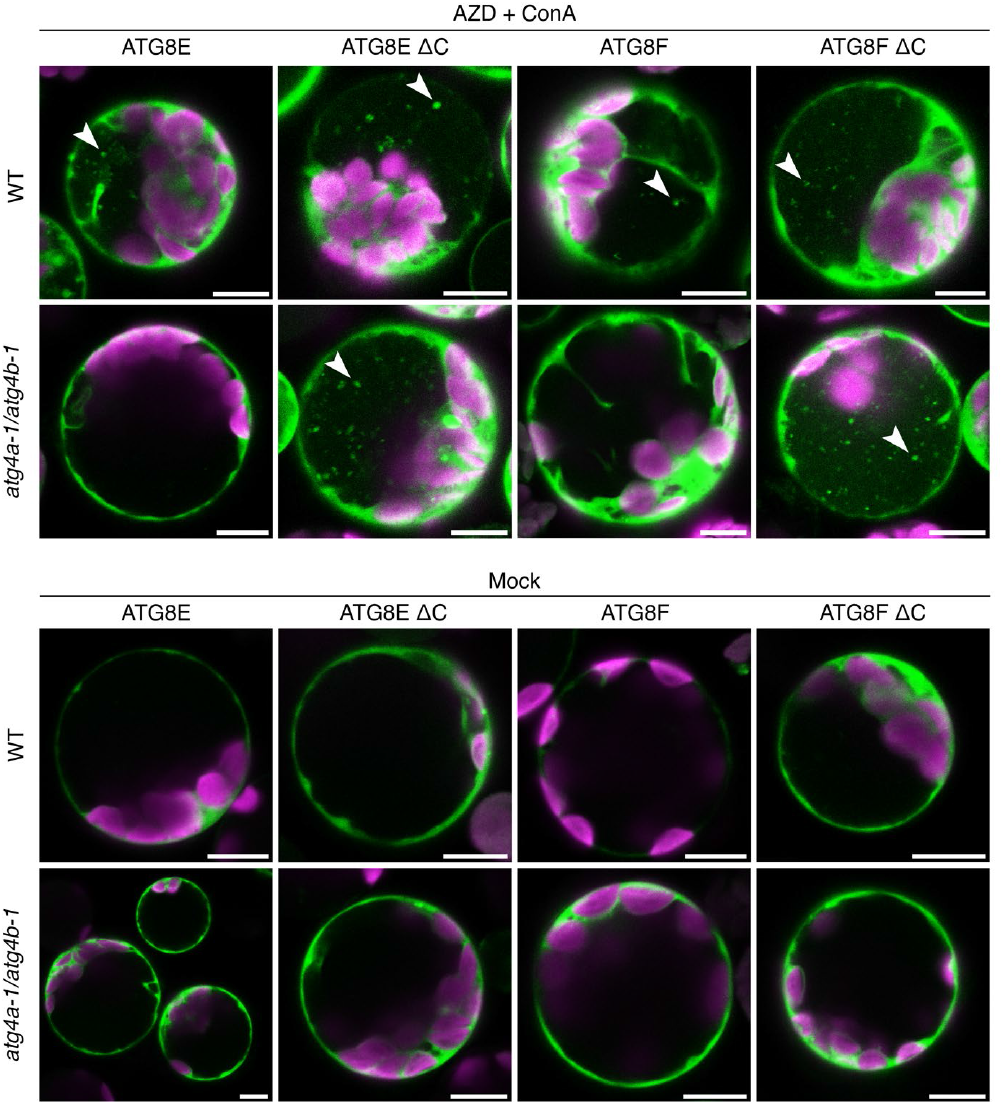
Artificially primed ATG8E and ATG8F restore accumulation of autophagic bodies in *atg4a-1/b-1*. Confocal images of mesophyll protoplasts isolated from 3-week-old WT and *atg4a-1/4b-1* (both in Wassilewskija-0 ecotype) plants. Protoplasts were transformed to transiently express GFP fusions of full-length and artificially primed versions of ATG8E and ATG8F isoforms. Protoplasts were either kept under standard conditions (mock) or subjected to 24 h treatment with AZD/ConA prior to imaging. White arrowheads point at autophagic bodies accumulating in the vacuoles of cells expressing primed versions of ATG8E and ATG8F when treated with drugs to induce autophagy. Scale bars, 10 μm.

**Figure S6.**
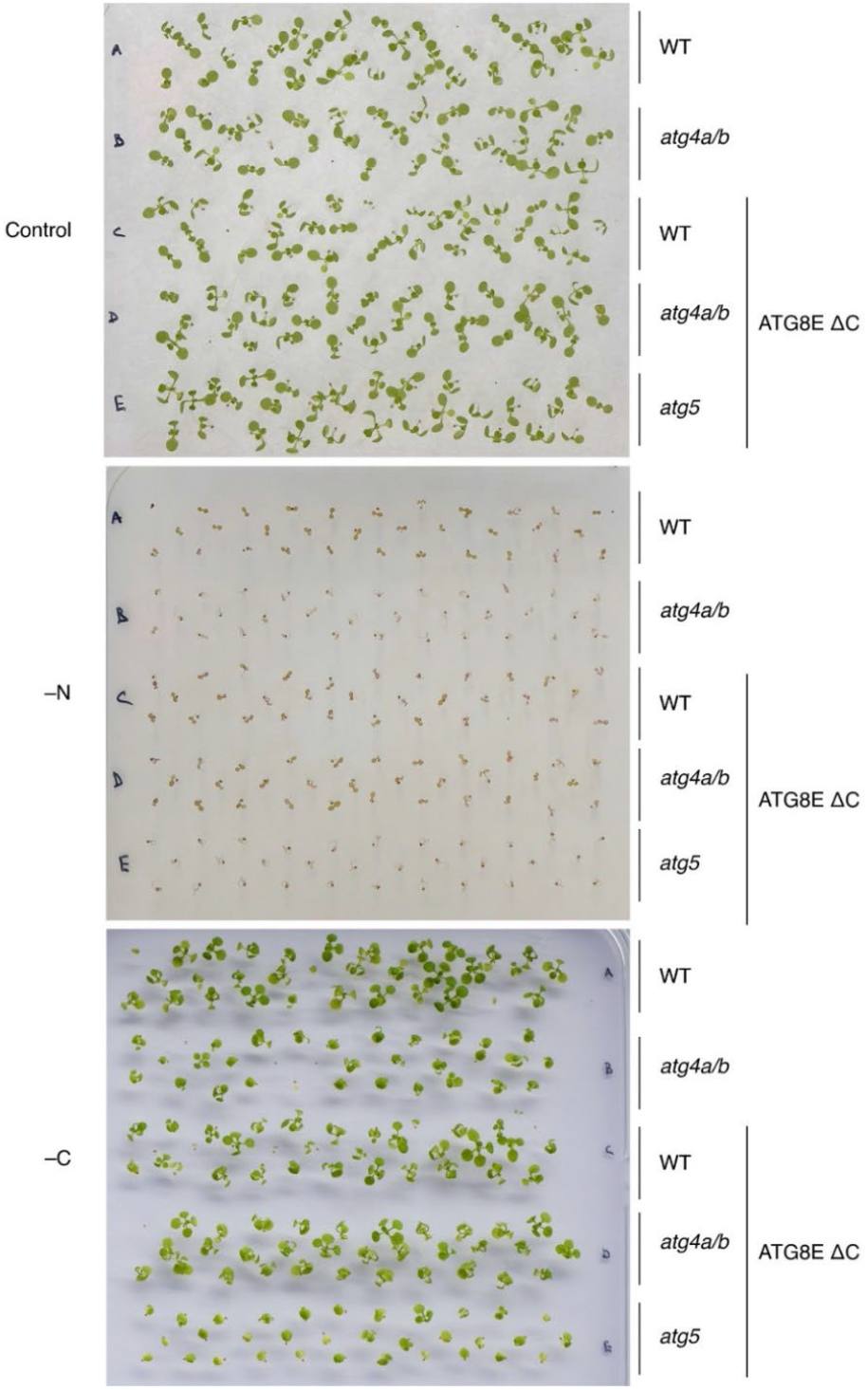
Expression of ATG8E ΔC complements autophagy-deficient shoot phenotypes of *atg4a/b*. Images of Petri plates with Arabidopsis seedlings grown on 0.5x MS medium (Control), nitrogen-depleted (–N) medium or carbon-depleted (–C) medium. Autophagy-deficient seedlings have decreased viability under –N conditions (quantification shown **Fig. 3E**). However, *atg4a/b* seedling expressing ATG8E **Δ**C exhibit WT-like viability. Seedlings grown on –C medium were subjected to 7 days of dark treatment followed by seven days of recovery prior to imaging. Autophagy-deficient seedlings are slower to recover compared to both wild-type seedlings and *atg4a/b* seedlings expressing ATG8E **Δ**C (quantification shown in **Fig. 3F**).

**Figure S7.**
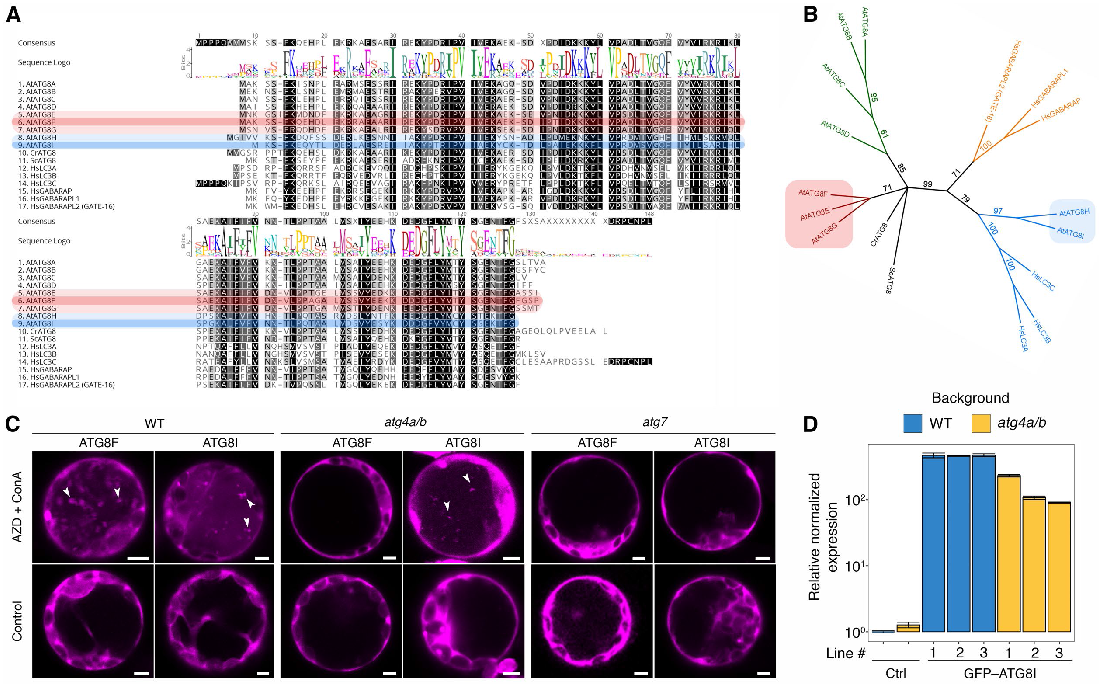
Transient overexpression of the natively truncated ATG8I isoform restores accumulation of autophagic bodies in the vacuoles of *atg4a/b* cells. **A**. Clustal W protein sequence alignment of *Arabidopsis thaliana* (At), *Chlamydomonas reinhardtii* (Cr), *Saccharomyces cerevisiae* (Sc) and *Homo sapiens* (Hs) orthologs of ATG8. Sequences of complete and truncated versions of Arabidopsis ATG8s with the highest expression levels, ATG8F and ATG8I, are highlighted in red and blue, respectively. Sequences of their closest paralogs are highlighted in lighter shades of the same colors. **B**. Phylogenetic tree illustrating evolutionary distances of ATG8 proteins compared in **A**, shows the separation of Arabidopsis ATG8s into the previously described two clades^14^: Clade I, more similar to fungi and comprising only complete ATG8s containing the C-terminus, and Clade II, comprising natively truncated ATG8s, more similar to animal ATG8s. **C**. mScarlet–ATG8F or mScarlet–ATG8I were transiently expressed in mesophyll protoplasts of WT, *atg4a/b*, or *atg7* Arabidopsis plants. An aliquot of each transformed protoplast suspension was subjected to AZD/ConA treatment for 17 h prior to imaging. Scale bars, 5 μm. White arrowheads indicate autophagic bodies. **D**. qPCR analysis of ATG8I expression in WT and *atg4a/b* lines with and without additional transgenes. Expression was normalized to the ATG8I level detected in WT and to three reference genes. Error bars show SEM for the technical triplicates. Three individual lines were checked for each genotype.

**Figure S8.**
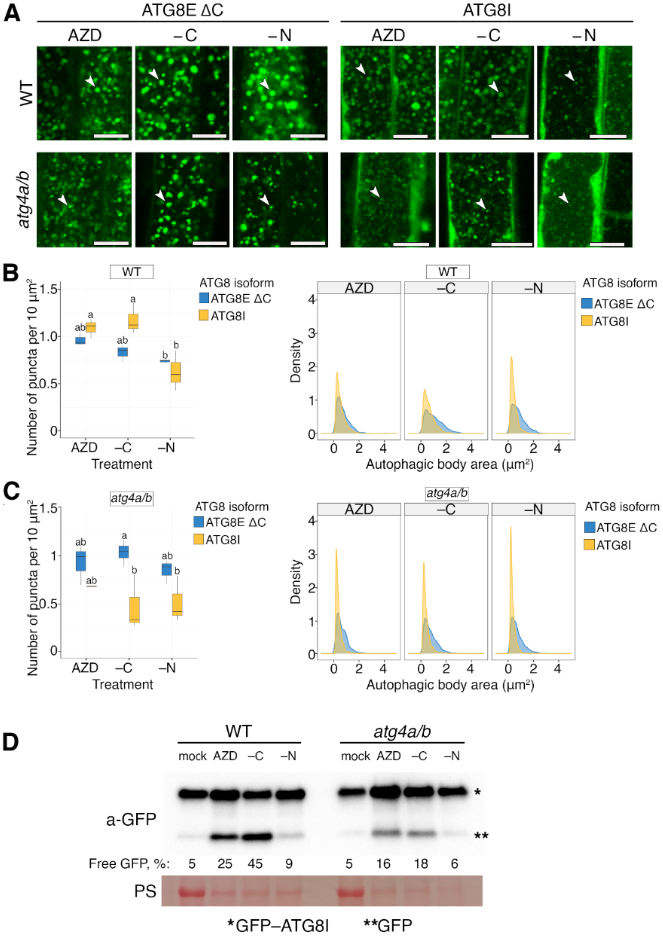
Overexpression of the natively truncated ATG8I isoform in WT leads to a decreased autophagic activity under nitrogen-depleted conditions. **A**. Confocal microscopy images showing phenotypes of autophagic bodies in the vacuoles of root epidermal cells of seedlings expressing ATG8E ΔC or ATG8I in WT and *atg4a/b* backgrounds. Seven-day-old Arabidopsis seedlings were subjected to treatments for 24 h prior to imaging. The treatments included AZD/ConA, –C/ConA, and –N/ConA. Scale bars, 10 μm. White arrowheads point at autophagic bodies in the vacuoles. **B**. Quantification of autophagic bodies number (left, box plot) and their size (right, density plot) for the WT seedlings illustrated in (**A**). Overexpression of ATG8I in the WT background led to a decreased number of autophagic bodies only under nitrogen-depleted conditions. Furthermore, ATG8I-overexpressing WT seedlings showed the most prominent decrease in the autophagic body size under low nitrogen conditions. Boxes indicate interquartile range (IQR); median is indicated by a horizontal line; whiskers represent range within 1.5 IQR; outliers are represented by empty circles. Tukey’s HSD test (α= 0.05), *n* = 18 biological replicates (510 technical replicates). The kernel density plot shows probability density function for autophagic body size depending on the expressed ATG8 isoform and treatment, *n* = 8001. **C**. Quantification analogous to the one shown in B performed for the *atg4a/b* seedlings. Tukey’s HSD test, *n* = 18 biological replicates (530 technical replicates). Kernel density plot, *n* = 9153. Overexpression of ATG8I in the *atg4a/b* background revealed lower number of autophagic bodies under all three conditions, indicative of the ATG8I isoform being insufficient for complete restoration of autophagic flux in the absence of other ATG8 isoforms. In agreement with this observation, ATG8I-overexpressing *atga4a/b* seedlings exhibited drastic decrease in autophagic body size under nitrogen-inducing conditions. **D**. GFP cleavage assay performed on the seedlings overexpressing GFP–ATG8I and illustrated in (**A**). The western blot confirms autophagic activity across all three stress conditions employed in the experiment, validates that nitrogen depletion produces the lowest autophagic flux when GFP–ATG8I is overexpressed and supports our observation that ATG8I fails to restore normal autophagic activity in *atg4a/b* pants. Densitometry results are shown as numbers under corresponding lanes and represent integrated density for the free GFP protein band expressed as % of total signal detected in the corresponding sample. PS, Ponceau S was used as a loading control.

**Figure S9.**
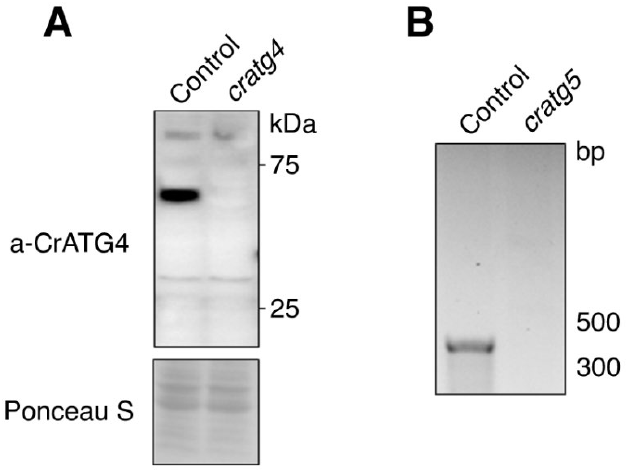
Validation of generated *Chlamydomonas ATG4* and *ATG5* knockout mutants. **A**. Western blot detection of ATG4 protein in the total protein extracts of WT (Control) and *ATG4* knockout (*cratg4*) *Chlamydomonas* cells confirms absence of ATG4 protein in the knockout background. Ponceau S staining was used as a loading control. **B**. Genotyping of the *ATG5* knockout mutant confirms deletion of the *ATG5* gene.

**Movie S1.**
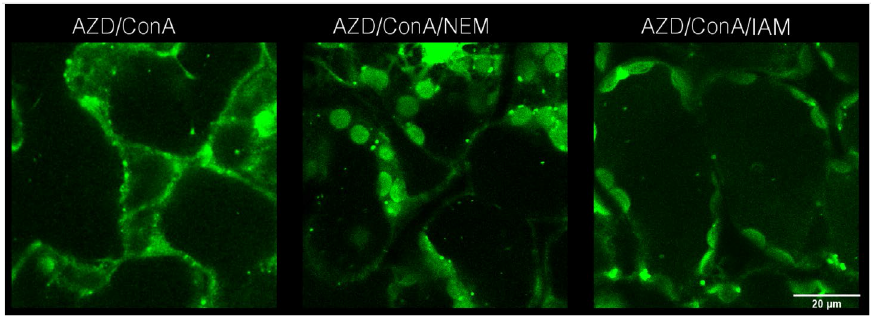
Chemical inhibition of ATG4 activity with IAM and NEM does not prevent accumulation of autophagic bodies in true leaves. Representative time-lapse scans of mesophyll cells of Arabidopsis true leaves expressing EosFP–ATG8E ΔC in the WT background. Leaves of one-month-old plants were infiltrated with MQ water containing 5 μM AZD8055 and 0.5 μM ConA additionally supplemented with 1mM IAM (Iodoacetamide) or 10 mM NEM (N-ethylmaleimide) without detaching them from plants. Leaves were imaged using CLSM 24 h after the treatment start. The video is available: https://youtu.be/vwG8cwUKy00

**Movie S2.**
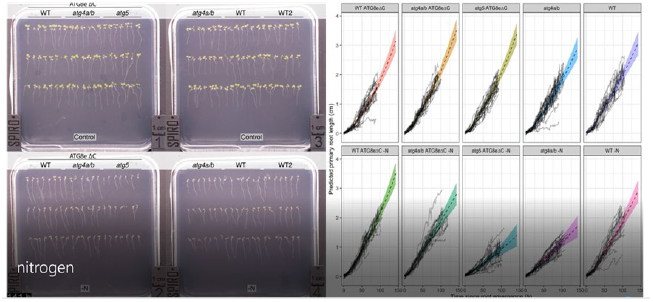
ATG8E ΔC complements stunted root growth phenotype of *atg4a/b* under nitrogen-depleted conditions. Representative SPIRO time-lapse data showing seedlings on control medium (top two plates) and–N medium (bottom two plates) imaged simultaneously. Charts on the right show data from the SPIRO root growth tracking assay plotted for each seedling and combined by genotype/growth conditions: black solid lines indicate root lengths plotted vs time for each seedling, dotted black lines show the predicted root length for each analyzed group, colored area indicates standard error for the root length prediction. Autophagy-deficient genotypes (*atg4a/b* and *atg5* ATG8E ΔC) show stunted growth under –N conditions, while *atg4a/b* ATG8E ΔC seedlings have normal root elongation under the same conditions. The video is available: https://youtu.be/jK_q8EZGVOE

**Movie S3.**
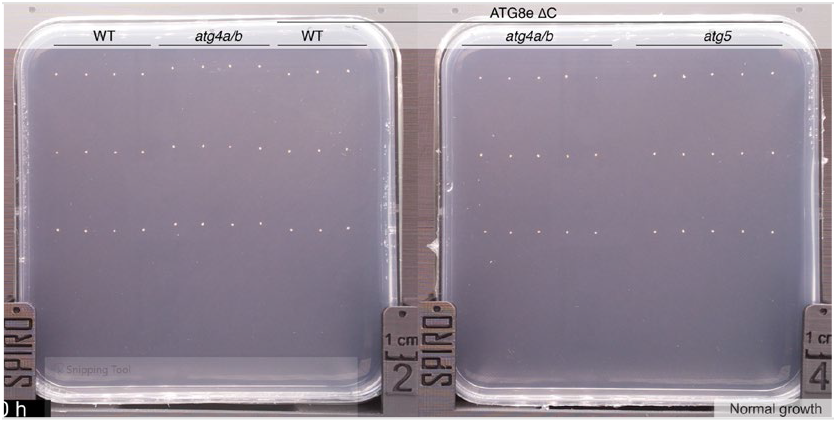
ATG8E ΔC complements weak root growth recovery of *atg4a/b* after carbon-depletion. Representative SPIRO time-lapse data showing seedlings on carbon-depleted plates imaged simultaneously. Seeds were allowed to germinate, and seedlings were grown under normal growth conditions, after which the lights in the growth cabinet were turned off and imaging proceeded for 4 days in the dark (carbon depletion), followed by imaging for another four days under normal growth conditions (recovery). Roots of autophagy-deficient seedlings (*atg4a/b* and *atg5* ATG8E ΔC) do not restart growth during recovery stage, while roots of *atg4a/b* ATG8E ΔC seedlings elongate normally under the same conditions. The video is available: https://youtu.be/VweCLd5lw0g

#### Pdf versions of main figures

##### Supplementary tables

**Table S1.**
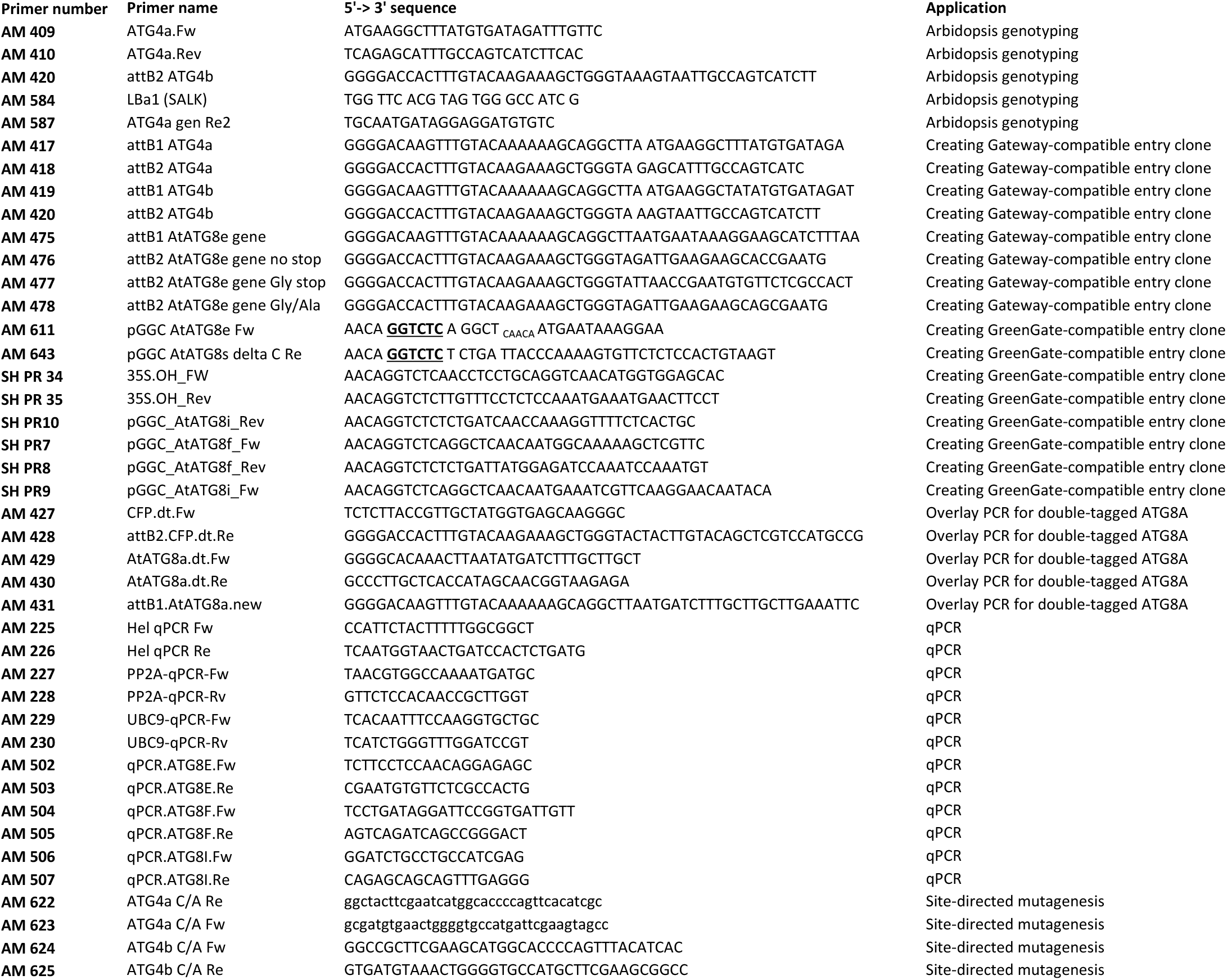
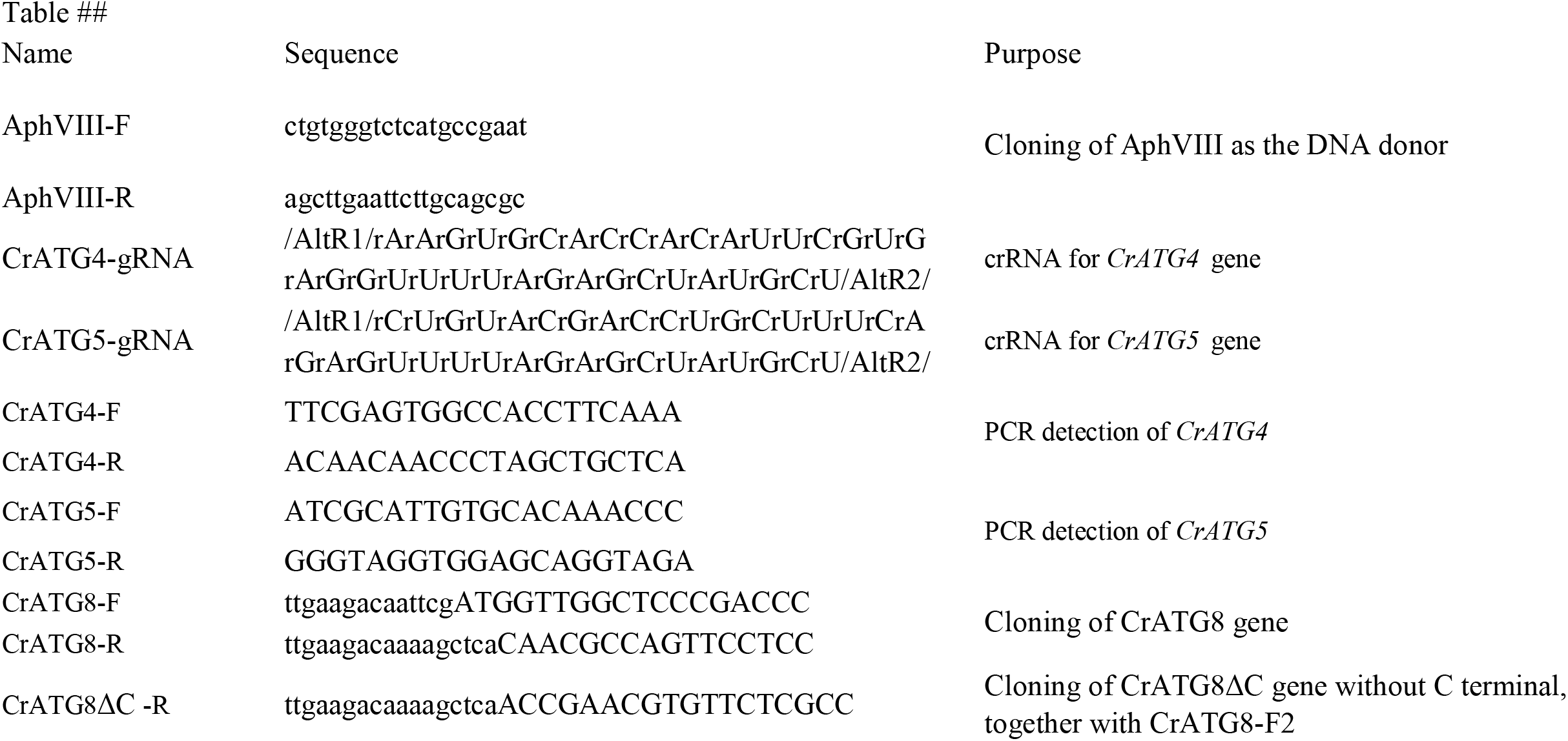
Primers used in this study.

**Table S2.** Plasmids used in this study.

| Construct number | Construct name | Insert | Selection marker in | Selection marker in | Vector type |
| --- | --- | --- | --- | --- | --- |
|  |  |  | plants | bacteria |  |
| AM 779 | pGGZ 2x35S::E2GFP-AtATG8E | GFP-ATG8E CDS | FastRed | spec | Expression in plants |
| AM 556 | pDONR/Zeo AtATG8A gene-CFP | AtATG8A gene-CFP | - | zeo | Entry clone |
| AM 567 | pUBN ubi10::EOSFP-AtATG8A gene-CFP | AtATG8A gene-CFP | basta | spec | Expression in plants |
| AM 655 | pDONR/Zeo AtATG8E gene | AtATG8E gene | - | zeo | Entry clone for Gateway cloning |
| AM 656 | pDONR/Zeo AtATG8E gene Gly stop | AtATG8E gene Gly stop | - | zeo | Entry clone for Gateway cloning |
| AM 657 | pDONR/Zeo AtATG8E gene Gly/Ala | AtATG8E gene Gly/Ala | - | zeo | Entry clone for Gateway cloning |
| AM 661 | pUBN ubi10::EOSFP-AtATG8E gene no stop | AtATG8E gene no stop | basta | spec | Expression in plants |
| AM 662 | pUBN ubi10::EOSFP-AtATG8E gene Gly stop | AtATG8E gene Gly stop (ATG8E delta C) | basta | spec | Expression in plants |
| AM 663 | pUBN ubi10::EOSFP-AtATG8E gene Gly/Ala | AtATG8E gene Gly/Ala | basta | spec | Expression in plants |
| AM 753 | pDONR/Zeo AtATG4A CDS | AtATG4A CDS | - | zeo | Entry clone for Gateway cloning |
| AM 754 | pDONR/Zeo AtATG4B CDS | AtATG4B CDS | - | zeo | Entry clone for Gateway cloning |
| AM 755 | pDONR AtATG4A C/A | AtATG4A CDS C/A mutant | - | zeo | Entry clone for Gateway cloning |
| AM 760 | pDONR AtATG4B C/A | AtATG4B CDS C/A mutant | - | zeo | Entry clone for Gateway cloning |
| AM 780 | pGGZ 2x35S::E2GFP-AtATG8F | GFP-ATG8F CDS | FastRed | spec | Expression in plants |
| AM 783 | pGGZ 2x35S::E2GFP-AtATG8I | GFP-ATG8I CDS | FastRed | spec | Expression in plants |
| AM 810 | pDEST15 GST-AtATG4A WT | GST-ATG4A WT | - | amp | Expression in bacteria |
| AM 811 | pDEST15 GST-AtATG4A C/A | GST-ATG4A C/A | - | amp | Expression in bacteria |
| AM 812 | pDEST15 GST-AtATG4B WT | GST-ATG4B WT | - | amp | Expression in bacteria |
| AM 813 | pDEST15 GST-AtATG4B C/A | GST-ATG4A C/A | - | amp | Expression in bacteria |
| AM 821 | pGGC AtATG8E delta C | AtATG8E delta C CDS |  | amp | Entry clone for GreenGate cloning |
| AM 822 | pGGC AtATG8F delta C | AtATG8F delta C CDS |  | amp | Entry clone for GreenGate cloning |
| AM 831 | pGGZ 2x35S::E2GFP-AtATG8E delta C | GFP-ATG8E delta C CDS | FastRed | spec | Expression in plants |
| AM 832 | pGGZ 2x35S::E2GFP-AtATG8F delta C | GFP-ATG8F delta C CDS | FastRed | spec | Expression in plants |
| SH 117 | pGGA 2x35S | 2x35S |  | amp | Entry clone for GreenGate cloning |
| SH 17 | pGGZ APA1 pr FLuc-mScarlet AtATG8F/RLuc:sGFP:NLS/kanR | FLuc-mScarlet AtATG8F | Kan | spec | Expression in plants |
| SH 19 | pGGZ APA1 pr FLuc-mScarlet AtATG8I/RLuc:sGFP:NLS/kanR | FLuc-mScarlet AtATG8I | Kan | spec | Expression in plants |
| SH 3 | pGGC AtATG8F | AtATG8F CDS |  | amp | Entry clone for GreenGate cloning |
| SH 4 | pGGC AtATG8I | AtATG8I CDS |  | amp | Entry clone for GreenGate cloning |

**Table S2.**
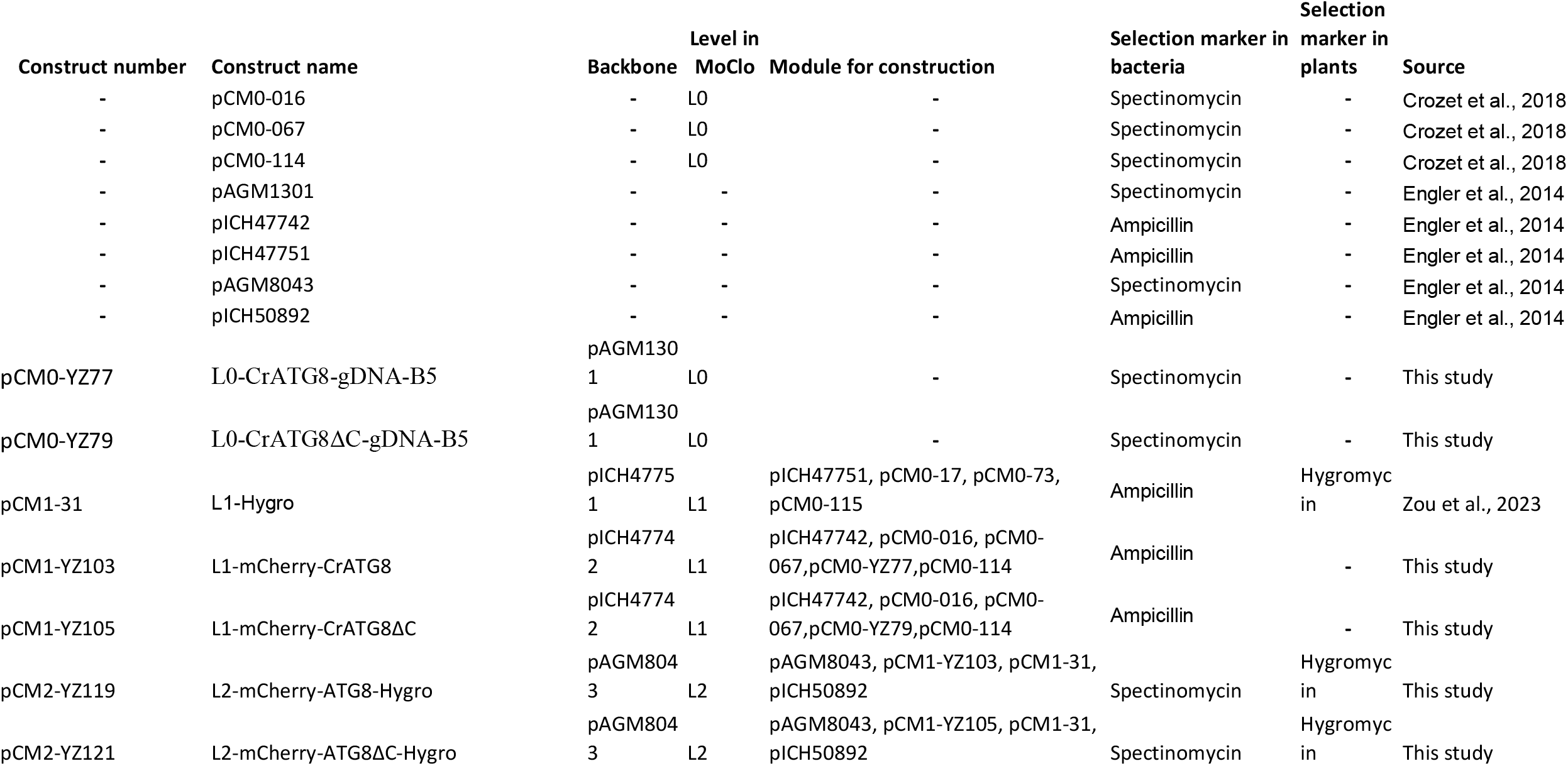
Plasmids used in this study.

**Table S3.** Arabidopsis transgenic lines used in this study.

| Seed batch | Transgene | Background | Comment |
| --- | --- | --- | --- |
| AM A 287-288 | 2x35S::E2GFP-ATG8F | WT | Good expression |
| AM A 289-290 | 2x35S::E2GFP-ATG8I | WT | Good expression |
| AM A 293-295 | 2x35S::E2GFP-ATG8F | atg4a-2/b-2 | Good expression |
| AM A 296-298 | 2x35S::E2GFP-ATG8I | atg4a-2/b-2 | Good expression |
| AM A 299-301 | 2x35S::E2GFP-ATG8F | atg7 | Good expression |
| AM A 302 | 2x35S::E2GFP-ATG8I | atg7 | Good expression |
| AM K 731-733 | pUBN ubi10::EosFP-ATG8E | WT | Somatic silencing |
| AM K 734-736 | pUBN ubi10::EosFP-ATG8E ΔC | WT | Good expression |
| AM K 737 | pUBN ubi10::EosFP-ATG8E G/A | WT | Somatic silencing |
| AM K 525 | pUBN ubi10::EosFP-ATG8E | atg4a-2/b-2 | Somatic silencing |
| AM K 526 | pUBN ubi10::EosFP-ATG8E ΔC | atg4a-2/b-2 | Good expression |
| AM K 527 | pUBN ubi10::EosFP-ATG8E G/A | atg4a-2/b-2 | Somatic silencing |
| AM K 819-820 | pUBN ubi10::EosFP-ATG8E ΔC | WT | Good expression |
| AM K 817-818 | pUBN ubi10::EosFP-ATG8E ΔC | atg4a-2/b-2 | Good expression |
| AM K 814-816 | pUBN ubi10::EosFP-ATG8E ΔC | atg5 | Ok expression |
| AM K 850 | pUBN ubi10::EosFP-ATG8A gene-CFP | WT | Good expression |
| AM K 851 | pUBN ubi10::EosFP-ATG8A gene-CFP | atg4a | Good expression |
| AM K 852 | pUBN ubi10::EosFP-ATG8A gene-CFP | atg4b | Good expression |
| AM K 853 | pUBN ubi10::EosFP-ATG8A gene-CFP | atg4a-2/b-2 | Good expression |
| AM K 507 | pMDC43 2x35S::GFP-ATG8E | WT | Somatic silencing |
| AM K 609 | pMDC43 2x35S::GFP-ATG8E ΔC | WT | Somatic silencing |
| AM K 612 | pMDC43 2x35S::GFP-ATG8E G/A | WT | Somatic silencing |
| AM K 648 | pMDC43 2x35S::GFP-ATG8E | atg4a-2/b-2 | Somatic silencing |
| AM K 649 | pMDC43 2x35S::GFP-ATG8E ΔC | atg4a-2/b-2 | Somatic silencing |
| AM K 650 | pMDC43 2x35S::GFP-ATG8E G/A | atg4a-2/b-2 | Somatic silencing |
| AM K 855 -856 | pUBN ubi10::EosFP-ATG8E ΔC | atg4a-2 | Good expression |
| AM K 857-858 | pUBN ubi10::EosFP-ATG8E ΔC | atg4b-2 | Good expression |
| AM A 433 (1-21) | 2x35S::E2GFP-ATG8F ΔC | WT | Good expression |
| AM A 425 (1-26) | 2x35S::E2GFP-ATG8F ΔC | atg4a-2/b-2 | Good expression |

## Notes

### Competing Interest Statement

The authors have declared no competing interest.

### Summary of Updates

Major revision of the original manuscript. The new version includes additional experiments supporting main conclusions.

